# An epigenomic investigation of atrial fibrillation in a matched left and right atrial human cohort

**DOI:** 10.1101/2025.08.29.673028

**Authors:** Adrian Rodriguez, Stephanie Frost, Alison M Thomas, Diego Fernandez-Aroca, Jishan Choudhury, Georgios Tikkas, Andrew Tinker, Patricia B Munroe, Christopher G Bell, Diego Villar

## Abstract

As the most prevalent cardiac arrythmia, atrial fibrillation is an important contributor to cardiovascular morbidity and mortality. Human population findings increasingly support its complex genetic architecture, with most genetic association signals for atrial fibrillation found in the non-coding genome. In this study, we integrated genome-wide histone modification, gene expression and DNA methylation levels in a paired left and right atrial cohort comprising permanent atrial fibrillation patients and sinus rhythm controls. First, we first identified epigenomic regions enriched in histone H3 lysine 27 acetylation (H3K27ac) across left and right atria from patients and controls, and associated them with differentially expressed genes to derive a set of dysregulated candidate loci – including *NPPB* and *SCX*. Second, by incorporating an independent replication cohort, we were able to validate gene expression and epigenomic differences for a subset of these candidate loci. Third, we profiled base-resolution DNA methylation levels and identified differentially-methylated regions (DMRs) between atrial fibrillation and sinus rhythm samples. Integration of these data with histone modification levels and gene expression allowed us to propose epigenetic mechanisms underlying transcriptomic and epigenomic changes across dysregulated loci, such as disruption of transcription factor binding by DNA methylation at the *LRRC4B* locus. The data and analyses we report constitute a systematic investigation of gene regulatory alterations across the left and right atria in permanent atrial fibrillation.

## INTRODUCTION

Atrial fibrillation (AF) is the most common form of cardiac arrythmia, affecting 0.5-1% of the worldwide population [1] and increasing up to 10% in the elderly. As such, AF significantly contributes to cardiovascular morbidity and mortality, and associates with other risk factors such as male gender, obesity, diabetes or smoking [2]. Mechanistically, AF is thought to arise from a complex interplay of triggers and substrate abnormalities that initiate and maintain arrythmia in the atria [3, 4]. Triggers are commonly caused by ectopic spontaneous firing in the pulmonary veins and can be maintained by re-entrant activity or rapidly discharging focuses. Subsequent remodeling of the atria - via altered activity of ion channels and structural changes favouring fibrosis – increases re-entry and can result in permanent AF [5, 6].

The genetic architecture of AF is complex and involves both rare and common variants that increase arrythmia susceptibility [7]. Approaches such as genetic linkage have attributed familial AF cases to mutations in genes coding for atrial natriuretic peptide[8], ion channel subunits [9–12], contractile proteins [13, 14] or cardiac transcription factors [15], among others – however, these mutations are absent in most early-onset AF cases. Genome-wide association studies (GWAS) have revealed over two hundred genetic loci associated to AF susceptibility [16–19]. Most of these association signals map to common genetic variation in the non-coding genome [20], with proposed effector genes including *PITX2* [21–23], *ZFHX3* [24, 25], HAND2 [21, 26, 27] or *SYNPO2L* [28, 29]. Collectively, AF GWAS loci are thought to act via effector genes leading to cardiac structural remodeling, as observed in atrial cardiomyopathy [7, 30].

Both the genetics and epidemiology of AF suggest an important role for gene regulation and epigenetic processes in mediating arrythmia susceptibility and cardiac remodeling [20, 31], which has prompted transcriptomic and epigenomic investigations in AF patient samples. First, previous studies have compared gene expression levels in human atria for AF patients and control individuals categorised as sinus rhythm (SR) (reviewed in [32]), with some interrogating paired right and left atrial samples [33–36]. Although observed gene expression differences across studies were often specific to patient cohorts, recent large-scale transcriptomic analyses found several putative AF susceptibility genes among those differentially expressed in the atria of AF patients [23]. Second, epigenomic studies of AF have identified genomic regions showing locus-specific variation in DNA methylation levels in AF [37–39], such as hypermethylation of the *PITX2* promoter in left atrium AF samples [40]. These analyses include epigenome-wide association studies (EWAS) of AF, albeit some have been conducted in blood rather than atrial tissue [38]. In contrast, AF epigenomic studies of other regulatory systems, such as histone modifications, have received less attention.

Here, we sought to integrate previously reported transcriptomic data in a paired left and right atrial appendage cohort [33] with epigenomic profiling of histone modifications and DNA methylation levels, obtained from a largely overlapping set of samples. By identifying candidate regions with differential epigenomic signals within the pathogenically critical tissue in AF patients and associating them with downstream gene expression, our results can inform the regulatory mechanisms underlying AF susceptibility and atrial remodeling.

## RESULTS

### An integrated epigenomic and transcriptomic dataset in left and right atrial appendages from atrial fibrillation patients and controls (related to Figure 1)

To investigate the regulatory landscape of human atria and its alterations in permanent AF, we conducted epigenomic profiling of histone modifications and DNA methylation in a set of paired left and right atrial appendage samples obtained from 7 atrial fibrillation (AF) patients and 5 sinus rhythm (SR) controls (Methods, Table S1, Figure S1a), and compared these data with previously obtained transcriptomes in a largely matched set of samples [33]. We first focused on genome-wide profiling of histone H3 lysine 27 acetylation (H3K27ac), a histone modification associated with active promoter and enhancer regions [41–43]. In line with previous reports [44–46], we identified an average of ∼50,000 enriched regions per sample, most of which were distal to annotated transcriptional start sites (Figure S1b-f).

**Figure 1:**
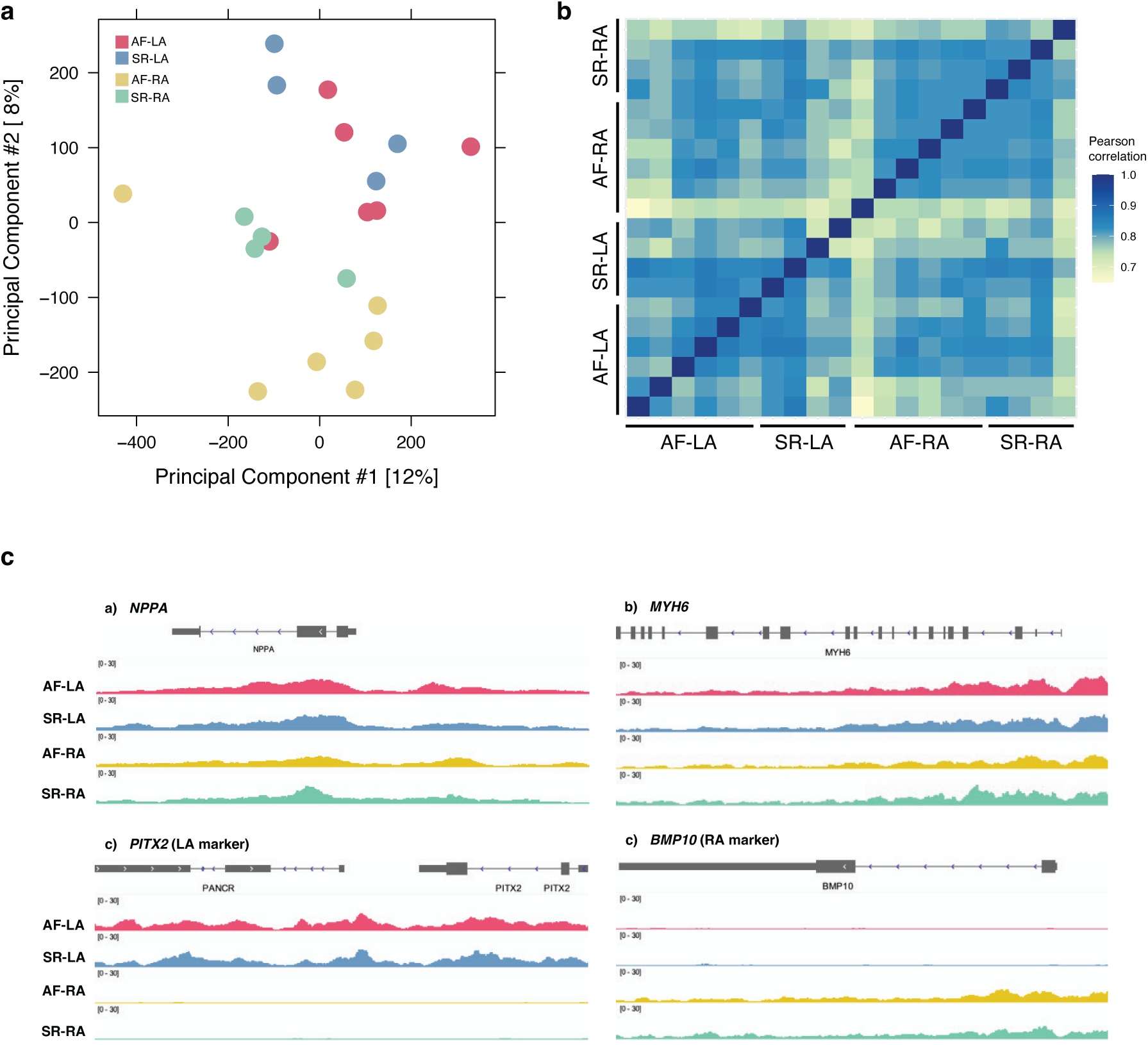
H3K27ac profiling across right- and left-atrial samples in a cohort of atrial fibrillation patients and sinus rhythm controls. **a.** H3K27ac enrichment differences across samples plotted on the first two principal components. Sample groups correspond to either left-(LA) or right-(RA) atrium samples from individuals with persistent atrial fibrillation (AF) or a normal heart rhythm (sinus rhythm, SR). Epigenomic differences more readily distinguish samples across anatomical side than disease status. **b.** H3K27ac enrichment levels across samples are represented as pairwise Pearson correlations. Samples corresponding to the same anatomical side tend to show higher correlations. **c.** Genome track examples of H3K27ac levels across the four sample groups for cardiac loci *MYH6* and *NPPA*, and *PITX2* and *BMP10* as left and right atrium loci, respectively. Each track represents average fold ChIP enrichment over input across samples in each group (Methods).

Principal components analysis of these data showed substantial heterogeneity in epigenomic signals across patients, with variance on the first two principal components mainly separating right atrium (RA) and left atrium (LA) samples (Figure 1A). Similarly, H3K27ac epigenomic enrichment showed higher pairwise correlations across samples from the same anatomical side, rather than disease status (Figure 1B). In agreement, examination of atrial expression markers associated to the left (*PITX2*) or right (*BMP10*) atrium showed clear differences in locus-level H3K27ac enrichment across samples from either anatomical side (Figure 1C). In contrast, cardiomyocyte loci such as *MYH6* and *NPPA* displayed similar enrichment levels in both right and left atrium samples. In sum, genome-wide H3K27ac profiling in atrial samples revealed widespread differences between anatomical sides, compared to more subtle epigenomic changes for disease status across samples.

### Definition of promoters and enhancers enriched in H3K27ac across disease and control samples (related to Figure 2)

We next aimed to identify promoters and enhancers showing an enrichment in H3K27ac levels between AF and SR samples in our data. To this end, we first used standard methods for differential analysis of ChIP-sequencing data [47, 48], comparing contrasts across anatomical side (H3K27ac differential enrichment in LA or RA) and disease status (AF or SR). Similarly to our previous results, we largely found differentially enriched regions between anatomical sides rather than disease status (TableS2), suggesting epigenomic differences across AF and SR samples may be subtle due to patient heterogeneity.

**Figure 2:**
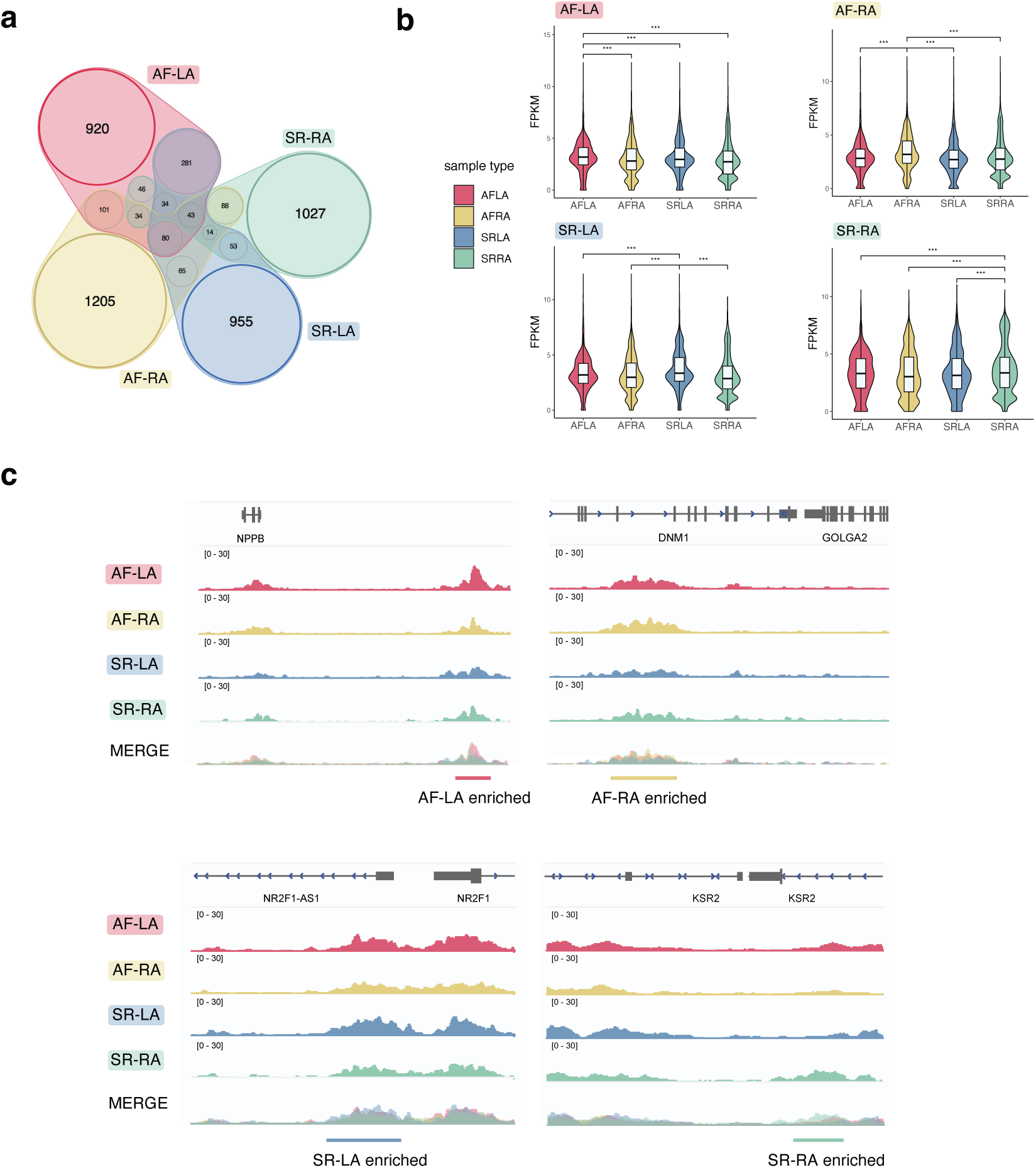
H3K27ac epigenomic regions enriched across sample groups. **a.** Venn diagram of epigenomic regions enriched in H3K27ac levels in each of the sample groups, as defined from pairwise comparisons in each category (Methods). Overlapping regions across sample groups were excluded from analysis (greyed colour shades). **b.** Violin plots of H3K27ac normalised read coverage (FPKM) over epigenomic regions enriched in each sample group (coloured labels, top left of each plot). Read coverage is represented in samples corresponding to each group (x-axis in each plot). As expected, H3K27ac coverage shows an upward shift in the sample group enriched regions were identified in. p-values correspond to one-sided Wilcoxon rank-sum tests with Bonferroni correction for multiple testing. ***: p < 0.001 **c.** Genome track examples of epigenomic regions enriched in each sample group, for *NPPB* (AF-LA), *DNM1/GOLGA2* (AF-RA), *NR2F1* (SR-LA) and *KSR2* (SR-RA) loci. Tracks represent average H3K27ac fold enrichment over input across samples in each group, and the overlay of signals across the four groups (Merge).

We therefore used an alternative approach to identify differential H3K27ac levels across AF and SR samples semi-quantitatively [48]. This strategy is based on (i) quantitative pairwise comparison of enrichment levels across samples to identify regions enriched in H3K27ac in each pair; and (ii) selection of enriched regions that are consistent across samples of the same type (Methods, Figure S2a-b). In this way, we defined ∼1,000 regions enriched in H3K27ac levels in each of the four sample groups (AF-RA, AF-LA, SR-RA and SR-LA) (Figure 2a; Table S3), or in AF or SR samples irrespective of anatomical side (Table S3). As expected for regions defined from H3K27ac signals, most were distal to transcriptional start sites (Figure S2c).

To confirm this approach recovers significant H3K27ac enrichments, we compared normalised read coverage for enriched regions in each sample group, and verified the expected upward shift in H3K27ac levels specifically for samples in the enriched group (e.g. AF-LA samples for AF-LA enriched regions, Figure 2b). We also compared average fold enrichment levels of H3K27ac for candidate enriched regions across sample groups (Figure 2c), and observed consistently higher histone mark levels in the enriched group. In agreement with the overall association between H3K27ac levels and gene expression [46, 49, 50], candidate loci identified in this manner include genes commonly upregulated in atrial fibrillation patients, such as *NPPB* [51]. By evaluating the intersection of enriched H3K27ac regions with gene ontology terms [52], we confirmed enriched regions in each sample group associate with cardiac biological processes (Figure S2d).

Lastly, we investigated the DNA sequences of H3K27ac enriched regions by measuring their overlap with transcription factor binding site motifs (TFBS) [53, 54], transcription factor (TF) binding locations from ChIP-sequencing catalogs [55] (Methods), and GWAS variants associated to cardiovascular traits [56]. These analyses were consistent with enriched epigenomic regions in each sample group showing an atrial regulatory signature, including sequence motifs and binding events of cardiac transcription factors (Figure S3a-b). Both GWAS variants associated to AF and other cardiovascular diseases were over-represented in H3K27ac enriched regions (Figure S3c), suggesting these regions include genetic susceptibility loci for AF and some of its co-morbidities.

In sum, by implementing a pairwise comparison approach for epigenomic profiling data in each sample group, we defined sets of candidate regions showing H3K27ac enrichment – which we next integrate with a largely matched gene expression dataset [33].

### Association of H3K27ac enrichment and differential gene expression defines candidate loci for altered gene regulation in atrial fibrillation (related to Figure 3)

We next sought to associate H3K27ac enriched regions across sample groups with downstream gene expression by leveraging previously published transcriptome data in the same sample cohort [33] (Figure S1, Table S1). First, we re-analysed this data and evaluated differential expression across anatomical side, disease status or both (Methods, Figure S4a-c, Table S4). We then associated enriched H3K27ac regions in each sample group with genes (Methods), and measured their enrichment for upregulated genes in the respective group (Fisher’s exact test, Methods). Whether across AF and SR samples from each anatomical side (Figure 3a) or overall (Figure S4d), we found enriched H3K27ac regions are significantly overrepresented in the vicinity of upregulated genes for the corresponding sample group. This observation is consistent with the role of H3K27ac as an active histone modification associated with promoter and enhancer activity [42], and predictive of downstream gene expression levels [50]. Moreover, it allowed us to select candidate loci with hallmarks of gene dysregulation in atrial fibrillation patients, by focusing on the subset of loci with both enriched H3K27ac and upregulated gene expression in each sample group (Figure 3b, Figure S4e and Table S5).

**Figure 3:**
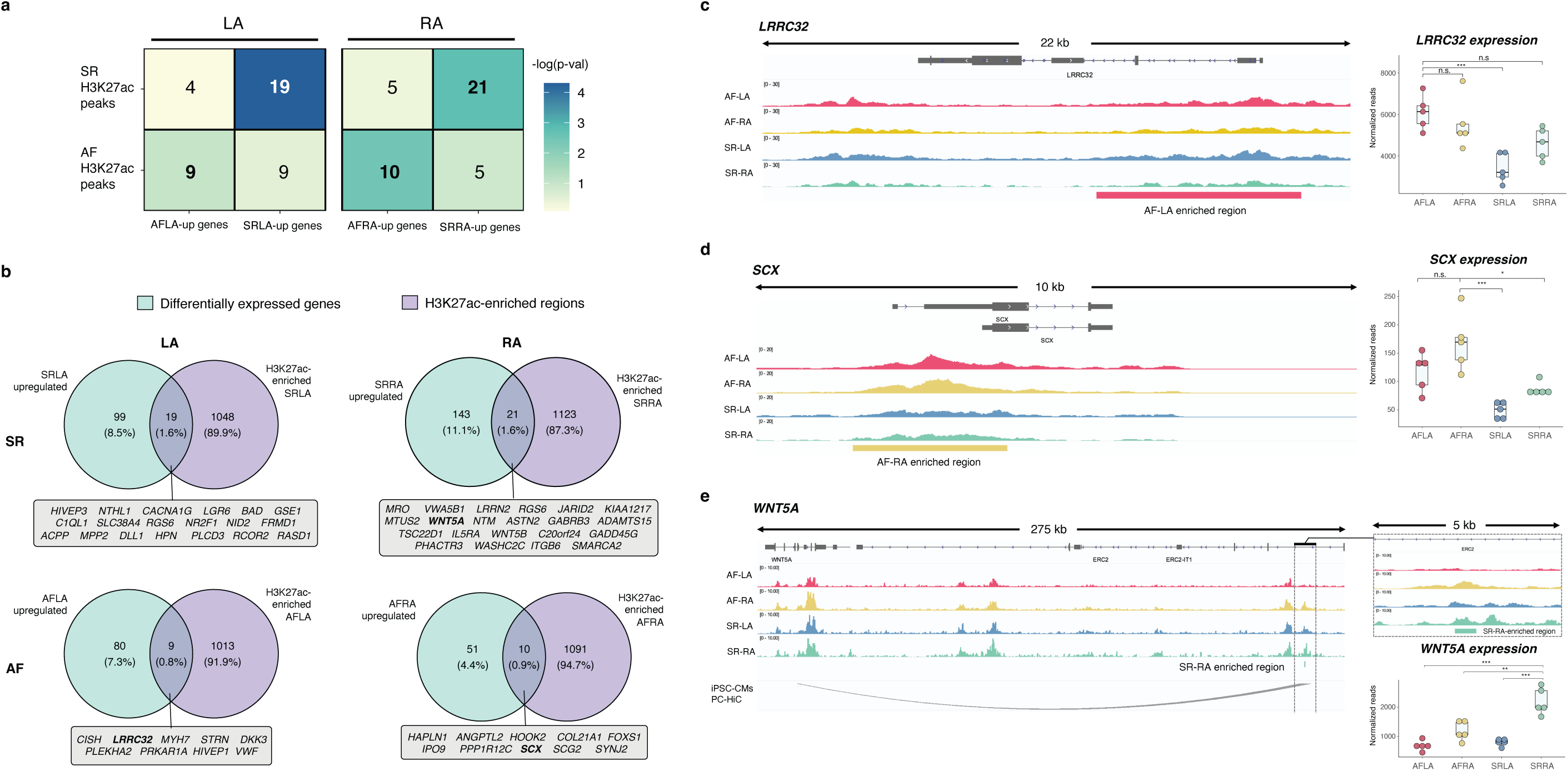
Association of H3K27ac enrichment with differential gene expression across sample groups and definition of candidate loci with altered gene regulation in atrial fibrillation. **a.** Association of differentially-expressed genes with enriched H3K27ac regions for each sample group. Numbers correspond to the number of gene loci with corresponding changes in gene expression (x-axis) and H3K27ac enrichment (y-axis). Cells are coloured by statistical significance (-log10 p-value, Fisher’s exact test). **b.** For each bolded comparison in a., Venn diagrams represent the overlap between differentially-expressed genes (cyan) and H3K27ac-enriched regions (violet). Overlapping gene loci (grey insets) correspond to candidate loci showing concordant gene expression and epigenomic changes in a particular sample group. **c.** An AF-LA enriched epigenomic region at the promoter region in the *LRRC32* locus (red bar), with gene expression levels being highest in this sample group (right handside, red points). n.s.: non-significant; ***: p < 0.001 (DESeq2 adjusted p-values). **d.** Example of an AF-RA enriched epigenomic region proximal to the *SCX* gene (yellow bar). Consistent with promoter activity, *SCX* expression levels are highest in AF-RA samples (right handside, yellow points). n.s.: non-significant; *: p < 0.05, ***: p < 0.001 (DESeq2 adjusted p-values). **e.** Example of a SR-RA enriched epigenomic region downstream of the *WNT5A* gene (green bar; right handside and top-right inset). External promoter-capture HiC data in iPSC-cardiomyocytes supports a 3D interaction between this putative enhancer region and the *WNT5A* promoter. *WNT5A* expression levels across sample groups are highest in SR-RA samples (bottom-right, light green). **: p < 0.01, ***: p < 0.001 (DESeq2’s adjusted p-values).

As illustrative examples, candidate loci with gene regulation changes in specific sample groups included *LRRC32* (AF-LA), *SCX* (AF-RA) and *WNT5A* (SR-RA) (Figure 3c-e). First, H3K27ac enriched regions in AF-LA samples included a segment spanning the promoter and part of the *LRRC32* gene body (Figure 3c). In agreement with this candidate region regulating proximal gene expression, we found *LRRC32* expression levels were highest in the AF-LA group, and significantly different from SR-LA samples (Figure 3c).

Second, in the *SCX* promoter region we identified an enriched H3K27ac element in AF-RA samples, suggesting increased promoter activity for *SCX* in this sample group. Accordingly, *SCX* gene expression levels were higher in AF, particularly for RA samples (Figure 3d). Third, we identified a distal region downstream of the *WNT5A* gene (in an intronic region of the *ERC2* gene, Figure 3e) with enriched H3K27ac levels in SR-RA samples. However, examination of previously published promoter-capture HiC data in iPSC-cardiomyocytes [57] supported an interaction of this element with the *WNT5A* promoter, suggesting potentially increased enhancer activity in SR-RA. Accordingly, *WNT5A* gene expression levels were increased in this sample group (Figure 3e, bottom right boxplot). Lastly, we similarly found consistent H3K27ac enrichment and gene expression levels for candidate loci with altered gene regulation in AF or SR samples (illustrative examples in Figure S4f-g).

On the whole, by focusing on the association between epigenomic H3K27ac levels and downstream gene expression, we defined a set of candidate loci with evidence of altered gene regulation in atrial fibrillation samples – which we next sought to validate in an independent replication cohort.

### Targeted validation of candidate loci in an independent patient cohort (related to Figures 4 and 5)

The candidate loci we identified here by integration of epigenomic and transcriptomic profiles were obtained in a relatively small atrial fibrillation cohort (Table S1, Figure S1, Methods). To evaluate the robustness of these candidates relative to donor biopsies, we obtained an independent set of 3 AF and 2 SR atrial samples to use as a replication cohort. As in our discovery cohort, these samples included matched RA and LA biopsies for each donor patient (Table S1, Figure S5a). To confirm their atrial origin and correct anatomical side assignments, we validated gene expression of cardiac, left atrium and right atrium markers in replication samples (Figure S5b).

**Figure 4:**
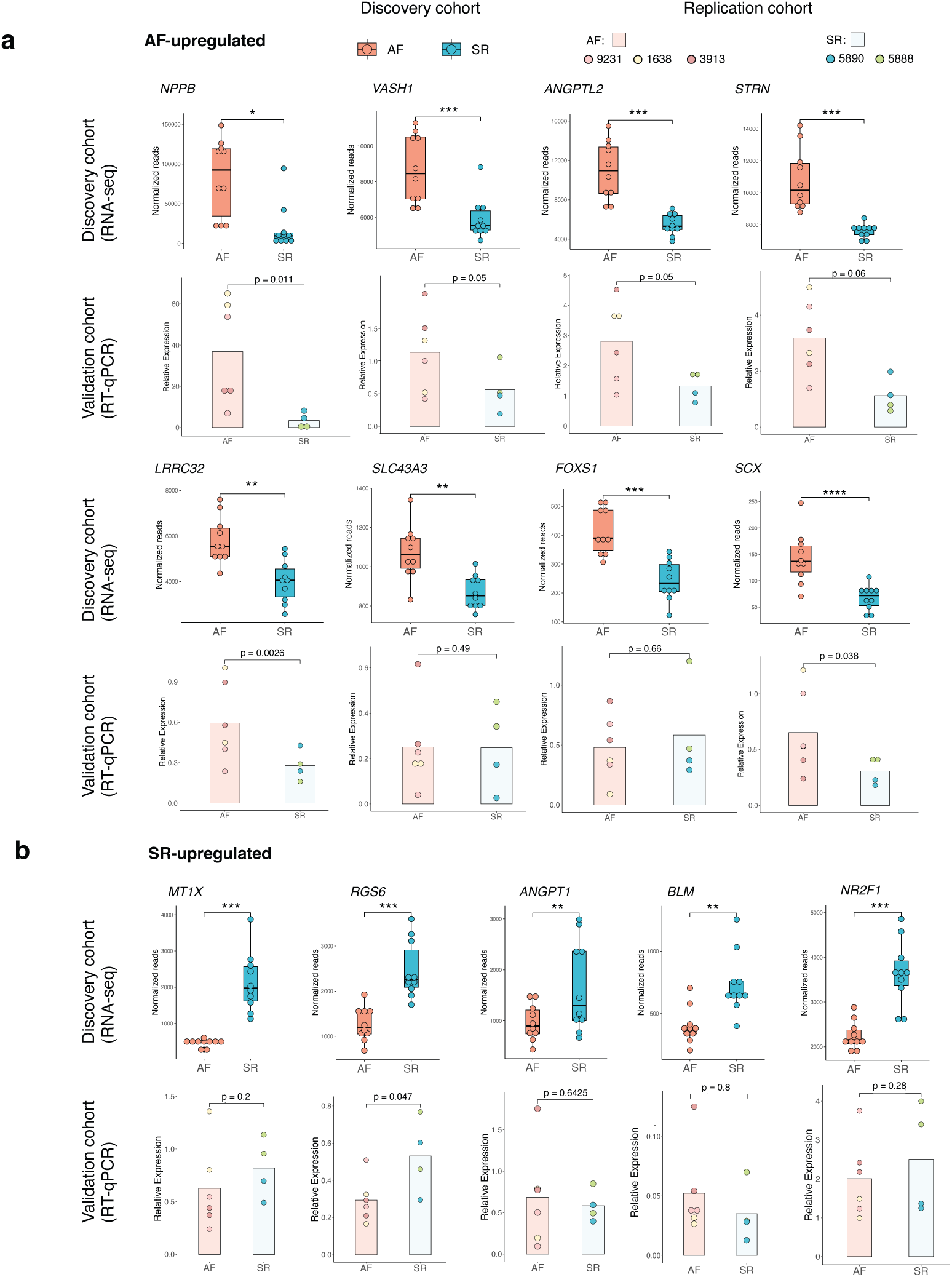
Targeted validation of gene expression levels at candidate loci in an independent replication cohort. Locus-specific comparison of gene expression levels in AF and SR samples for selected candidate loci in the discovery cohort (top boxplots in each category, RNA-seq), and their validation in an independent replication set (bottom barplots in each category, RT-QPCR). Gene expression differences across disease status were considered as validated in independent samples when the direction of change was maintained with a p-value < 0.2 (one-sided t-test). **a.** Gene expression levels in the discovery cohort (top boxplots) and validation cohort (bottom barplots) across 8 AF-enriched candidate loci. **b.** As in a. for five SR-enriched candidate loci. See also Figure S5 and Table S6.

**Figure 5:**
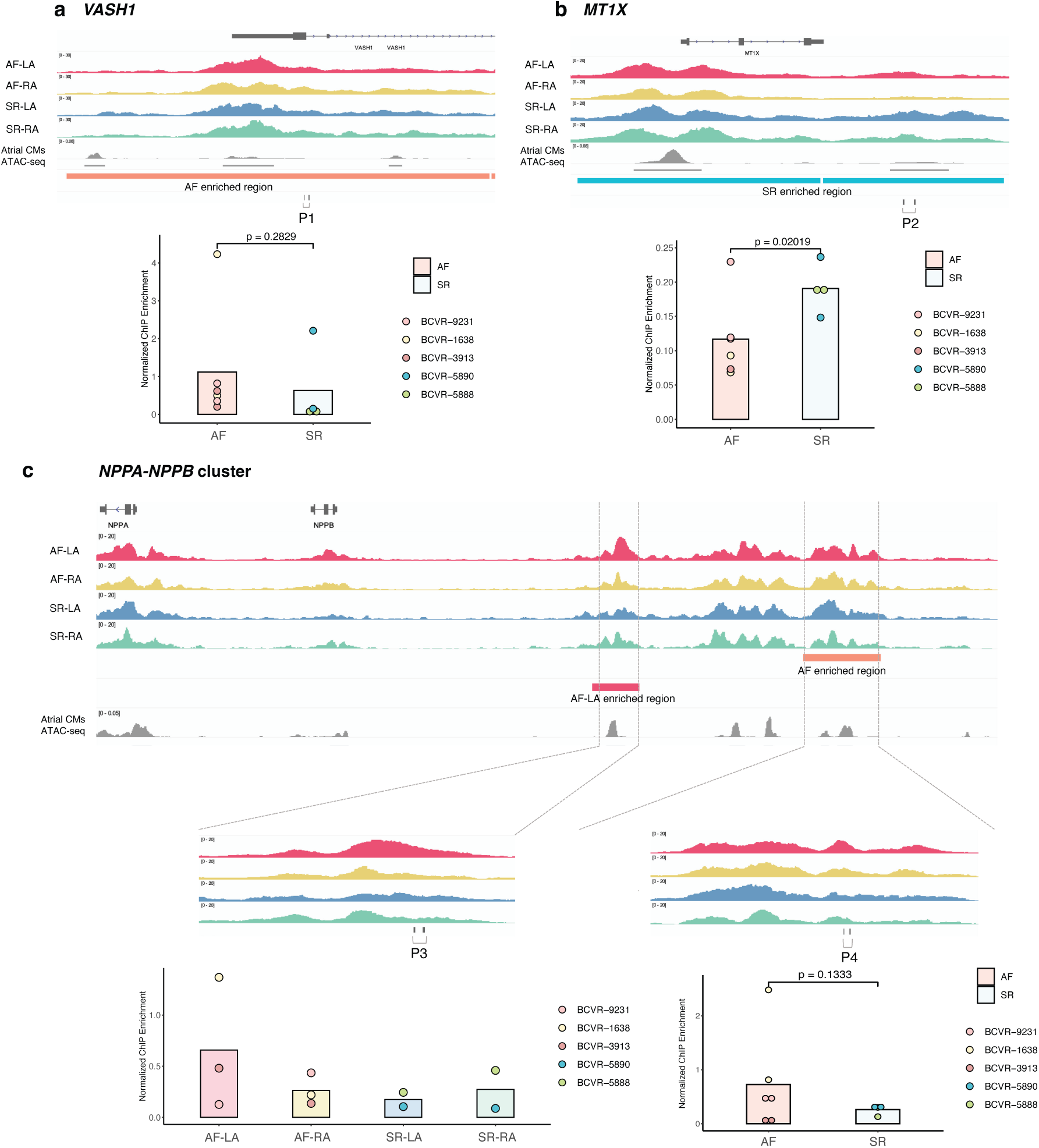
Targeted validation of H3K27ac enrichment at candidate loci in an independent replication cohort a –. **c.** Examples of ChIP-QPCR validation of H3K27ac enrichment for an AF-enriched region in the *VASH1* locus (**a**, sienna bar), a SR-enriched region in the *MTX1X* locus (**b**, cyan bar), and AF- and AF-LA enriched regions in the *NPPA/NPPB* locus (**c,** sienna and red bars, respectively). For each example, genome tracks show H3K27ac levels in each sample group (AF-RA, AF-LA, SR-RA, SR-LA), the location of validated enriched regions (coloured bars) and single-cell ATAC-seq signals corresponding to atrial cardiomyocytes in the human heart. P1 to P4 indicate primer pairs used for validation of each region (barplot insets). Highlighted p-values correspond to one-sided t-test. For the P3 region in c, ANOVA p-value = 0.54 See also Figure S5 and Table S6.

Towards validation of candidate loci, we used locus-specific quantitative PCR to measure gene expression levels (Figure 4) and H3K27ac enrichment (Figure S5) across a subset of candidate loci (Table S6). Specifically, we focused on evaluation of gene expression and histone mark enrichments across disease status (AF or SR samples) in the smaller replication cohort. For gene expression levels (Figure 4), we found an overall good correspondence for expression differences in the discovery and replication cohorts, with most candidate genes showing the expected expression shifts in the independent sample set. We validated gene expression differences in 9 out of 12 loci tested (validation rate ∼75%, p-value < 0.2 t-test), with similar rates for AF (Figure 4a, 6 out of 8 loci) or SR upregulation (Figure 4b, 3 out of 5).

In contrast, locus-specific H3K27ac enrichments in replication samples showed higher variability (Figure S5c-d), and a lower validation rate ∼50% (p < 0.2, t-test). This observation is consistent with the relatively large size of H3K27ac enriched regions compared to the genomic segments we tested (Methods, Figure S5). To illustrate this difference, we examined validation results by ChIP-QPCR in comparison with H3K27ac data in the discovery cohort across three candidate loci (Figure 5). First, for an AF-enriched H3K27ac region in the *VASH1* locus, we observed slightly higher H3K27ac levels by ChIP-QPCR in independent samples (Figure 5a, P1 primer pair, p = 0.28), with the tested region corresponding to a genomic segment proximal to an accessible region in human atrial cardiomyocytes [58]. Second, we validated differential H3K27ac levels for an SR-enriched region in the *MT1X* locus (Figure 5b). In this case, the region tested in the replication cohort overlapped an accessible region in human atrial cardiomyocytes (CMs) (P2, p = 0.02). Lastly, we examined validation results for two enriched regions in the *NPPB* locus, in AF-LA and AF samples, respectively. These map to a previously defined super-enhancer in mouse heart [59], and both tested regions overlapped accessible chromatin in atrial CMs (P3 and P4, Figure 5c). For either region, we observed the highest H3K27ac levels in the expected sample group, although differences were not significant for the AF-LA region.

Overall, locus-specific evaluation of gene expression and histone mark enrichments at a subset of our candidate loci in replication samples supported the majority of candidate loci can be validated in an independent cohort, lending robustness to the putative gene regulation alterations we identified in our discovery cohort.

### Base-resolution DNA methylation signals at candidate loci inform mechanisms of altered gene expression and TF binding (related to Figures 6 and 7)

To further investigate the mechanisms underlying altered gene regulation in atrial fibrillation patients for our candidate loci, we carried out genome-wide profiling of DNA methylation levels in 16 samples from our discovery cohort via the Illumina EPICv2 array (Methods, Table S1, Figure S1a). First, we performed an exploratory EWAS (Figure S6a, Table S7) for differentially-methylated cytosines (DMCs), and then a differentially-methylated region (DMR) analysis (Figure 6a) to identify regions with DNA methylation differences between AF and SR samples (Methods). As expected, assessment of single CpG methylation differences was underpowered in our sample set, and most signals we identified through EWAS (Table S7) did not overlap candidate loci. However, aggregating CpG methylation signals into DMRs resulted in a set of 148 regions discriminating AF and SR disease status for a majority of donors (Figure 6a-b).

**Figure 6:**
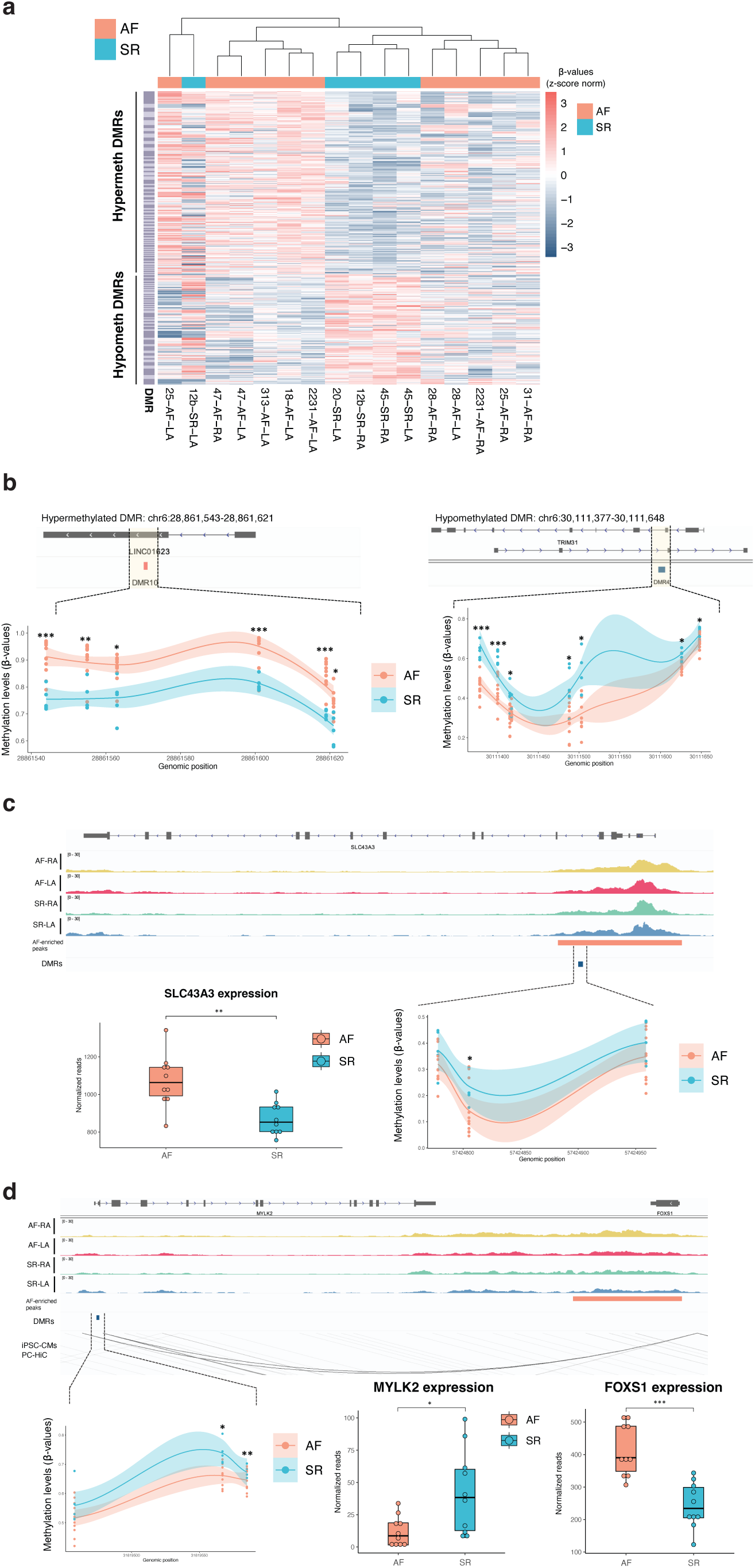
Differentially-methylated regions in AF samples inform gene dysregulation mechanisms in candidate loci. **a.** Heatmap of DMRs identified as hypermethylated (y-axis, top half) or hypomethylated (y-axis, bottom half) in AF samples. Heatmap scale corresponds to Z-score normalization of beta-values. Hierarchical clustering (x-axis, top; sienna for AF and cyan for SR) shows a subset of samples associates with DNA methylation differences across DMRs. **b.** Examples of DMRs hypermethylated (DMR10, left) and hypomethylated (DMR4, right) in AF patients. Shaded trendlines correspond to local regression (LOESS method) with confidence interval, and p-values to statistical significance of CpGs within a DMR. **c.** A hypomethylated DMR in *SLC43A3* associates with H3K27ac enrichment and increased expression levels in AF patients. Representation as in **a** and previous Figures. **d.** A hypomethylated DMR proximal to *MYLK2* acts as a potential enhancer of *FOXS1* expression and associates with H3K27ac enrichment and increased *FOXS1* expression in AF patients. Representation as in **a** and previous Figures. FDR-corrected two-sided t-test for differentially-methylated CpGs; ***: p < 0.001; **: p < 0.01; *: p < 0.05). DESeq2 adjusted p-values for gene expression differences: *: p < 0.05, **: p < 0.01; ***: p < 0.001 See also Table S7.

**Figure 7:**
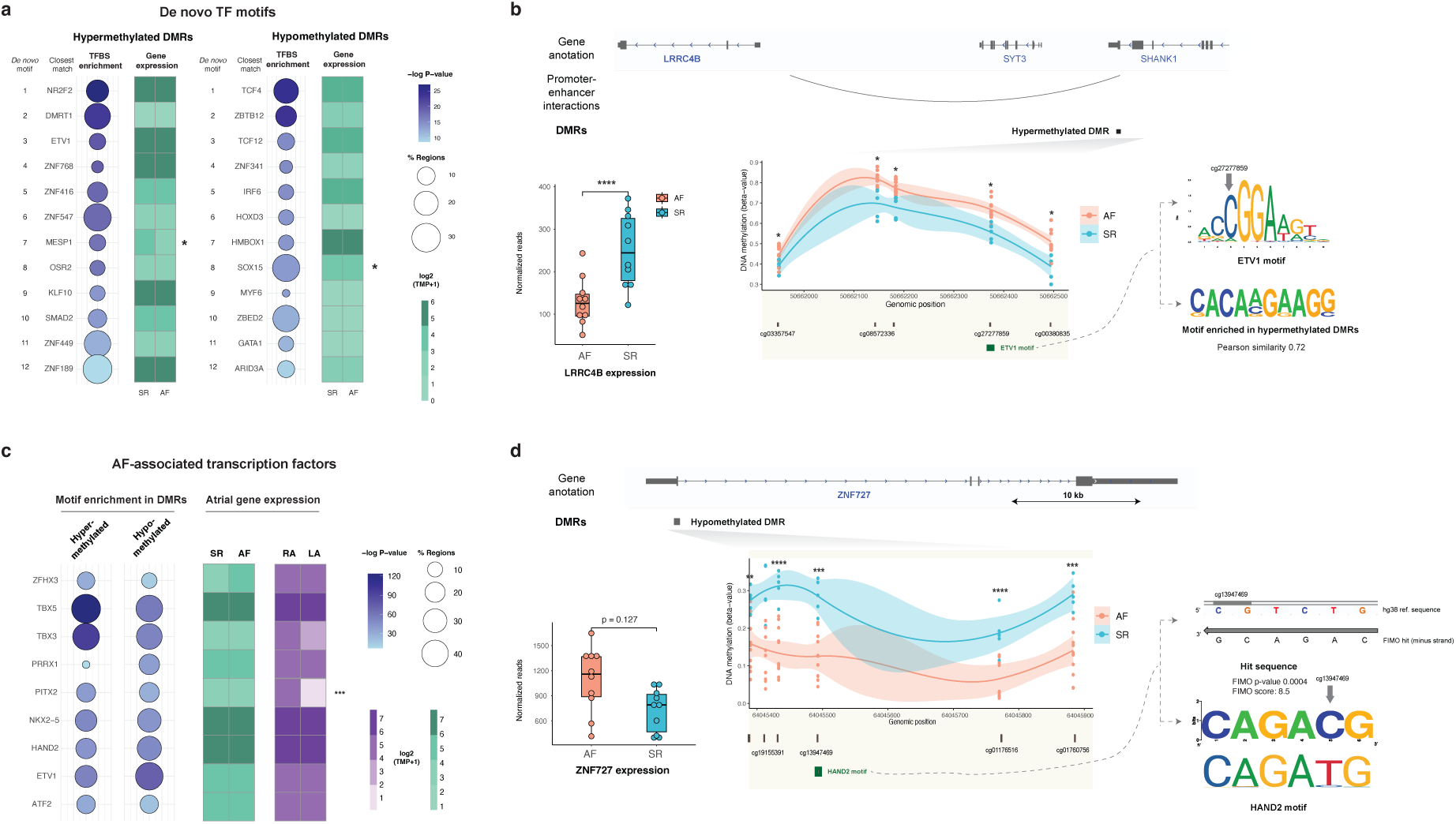
Transcription factor motif enrichment in AF differentially-methylated regions and examples of potential disruption of TF binding by DNA methylation. **a.** *De novo* transcription factor motifs enriched in hypermethylated and hypomethylated DMRs. Circle plots in each category indicate percentage of DMRs harbouring sequence motifs (circle size) and statistical significance (circle shade; −log10 p-value, hypergeometric test with Benjamini-Hochberg correction for multiple comparisons). Each enriched motif was assigned to the indicated transcription factors based on motif similarity and cardiac expression (Methods). Heatmaps indicate median expression levels (log2(TPM+1)) for each transcription factor in sinus rhythm controls (SR) or permanent AF patients (AF). Asterisks indicate differential gene expression (FDR p-value < 0.05; Methods). **b.** Example of a hypermethylated DMR in the *LRRC4B* locus (DMR26), for which one of the differentially-methylated CpGs overlaps an ETV1 motif. (Top tracks) Gene and DMR annotation, and promoter-enhancer interactions in this genomic locus. (Bottom middle inset) DNA methylation differences across the DMR, shown as beta-values for AF (sienna) and SR (cyan) samples. Annotations below the plot correspond to individual CpGs and the ETV1 sequence motif (green box). *: p-value < 0.05 (Bottom right inset) The ETV1 motif is highly alignable to the *de novo* motif enriched in hypermethylated DMRs (top-right comparison of motifs logos, Pearson correlation-based similarity score = 0.72; differentially methylated CpG cg27277859 marked on the top motif by grey arrow). (Bottom left boxplot) Expression levels of *LRRC4B* expression are significantly higher in SR patients (*: p < 0.05). Expression levels of other genes in this locus are similar in SR and AF samples (Figure S7b). **c.** (Circle plots, left) TFBS motif enrichment for AF-associated transcription factors in hypermethylated and hypomethylated DMRs. Representation and details as in **a.** (Heatmaps, right) Median gene expression levels of AF-associated transcription factors across SR or AF samples (shades of green); and right atrium (RA) or left atrium (LA) samples (shades of violet). Asterisks indicate differential gene expression (FDR p-value < 0.05; Methods). **d.** (Top-middle inset) Example of a hypomethylated DMR in the *ZNF727* locus, which includes a differentially-methylated CpG overlapping a sequence similar to the motif for AF-associated transcription factor HAND2 (FIMO score 8.5, p-value 0.0004). (Top tracks) Gene and DMR annotation. (Bottom middle inset) DNA methylation differences across the DMR. Annotations below the plot correspond to individual CpGs and the HAND2 sequence motif (green box). P-values: **: p < 0.01; ***: p < 0.001; ****: p < 0.0001. (Bottom left logo) Sequence logo for the HAND2 motif. The hypomethylated CpG cg13947469 is highlighted by a grey arrow. (Left boxplot) Expression levels of *ZNF727* are higher in AF patients (p=0.127). Details as in **b**. See also Figure S7 and Table S7.

To investigate DNA methylation differences across candidate loci, we next intersected DMRs with enriched H3K27ac regions (Figure 2, Figure S6b). As expected for CpG array data (Methods), DMRs were relatively proximal to annotated TSSs, were often outside CpG islands and showed limited overlap with enriched H3K27ac regions across sample groups (Figure S6b-d). We thus associated DMRs with candidate regions at the locus level, and explored the association of DNA methylation differences with differential gene expression and H3K27ac enrichment (Figure S6b and Table S8).

DNA methylation signals at candidate loci often informed potential mechanisms of altered gene regulation in atrial fibrillation. For the *SLC43A3* locus (Figure 6c), we identified a hypomethylated DMR (lower DNA methylation levels in AF) within an enriched H3K27ac promoter region in AF samples. Hypomethylation for this DMR associated with higher gene expression of *SLC43A3* in AF patients, as expected for increased promoter activity. In contrast, for the *FOXS1 / MYLK2* locus (Figure 6d), we identified a hypomethylated DMR in the *MYLK2* promoter region, and distally to an AF-enriched H3K27ac region with potential enhancer function. Expression levels of *MYLK2* were inconsistent with the identified DMR, and higher in SR controls compared to AF patients. However, promoter-capture HiC data [57] in this region supported an interaction between the AF-enriched H3K27ac region downstream (proximal to *FOXS1*) and the identified DMR, which may thus act as a distal enhancer of *FOXS1* expression. In agreement, *FOXS1* expression levels were found to be increased in AF patient samples. These examples illustrate how DNA methylation and H3K27ac levels can jointly inform epigenomic alterations at AF candidate loci.

We next sought to investigate the sequence composition of DMRs showing hyper- and hypomethylation in AF samples. To this end, we first identified TFBSs across DMRs, and observed distinct sets of TFBSs overrepresented in each group (Figure 7a, Figure S7, Methods). These include motif sequences for transcription factors previously associated to AF, such as ETV proteins [60]; and typically corresponded to transcription factors expressed at similar levels in AF and SR samples (Figure 7a). For instance, in the genomic locus spanning *LRRC4B, STY3* and *SHANK1* genes, we identified a hypermethylated DMR in which one of the CpGs overlapped an ETV-1 motif (Figure 7b). Gene expression levels of genes in this locus and promoter-capture HiC data in iPSC-cardiomyocytes [57] is consistent with this DMR acting as a distal enhancer of *LRRC4B*, a gene previously implicated in cardiac dysfunction and fibrosis [61, 62]. Integration of these data allowed us to hypothesise hypermethylation of the ETV-1 motif may result in lower activity of this enhancer in AF patients, as suggested by reduced *LRRC4B* expression (Figure 7b, Figure S7). In contrast, we found ETV-1 expression levels were similar in both sample groups (Figure 7a).

Lastly, we also associated DMRs with GWAS variants and transcription factor binding events [55, 56], and observed enriched biological processes and cardiovascular traits consistent with atrial regulatory mechanisms (Figure S7). In agreement, we observed TFBSs corresponding to AF-associated transcription factors (TFs) [19] were often also enriched in DMRs (Figure 7c), and these TFs were expressed in atrial samples with no clear specificity towards disease status or anatomical side (Figure 7c). Instead, we found instances where differences in DNA methylation occurred within the TFBS of an AF-associated TF (Figure 7d). Within a hypomethylated DMR in the *ZNF727* promoter (Figure 7c), we identified a sequence similar to the standard HAND2 motif for which hypomethylation in AF samples could facilitate HAND2-mediated expression. In agreement, we observed higher expression levels of *ZNF727* in AF patients.

In sum, DNA methylation changes associated to AF in our sample cohort often suggested altered mechanisms of gene regulation and TF binding underlie gene expression differences, thus providing testable hypotheses for targeted investigations.

## DISCUSSION

Human population studies increasingly support a complex genetic architecture for atrial fibrillation [3, 19], with most genetic association signals corresponding to non-coding regions with potential roles in gene regulation, such as tissue-specific enhancers [20, 63]. Accordingly, both transcriptomic and DNA methylation datasets support widespread gene regulation changes are associated with the development of persistent AF [38, 64]. Using a small set of matched left and right atrial biopsies comprising sinus rhythm controls and persistent AF patients, here we integrated epigenomic profiling of the histone modification H3K27ac and DNA methylation levels with transcriptomic data previously obtained in these samples [33]. This resource allowed us to identify epigenomic changes on histone modifications and DNA methylation in AF patients, and test their association with downstream gene expression levels. First, comparison of H3K27ac levels in each sample group with differential gene expression allowed us to define a set of candidate loci with altered gene regulation in AF, which enrich for GWAS loci associated to AF and its cardiovascular comorbidities. These candidates include both previous AF-associated loci such as *NPPB* and *ANGPTL2*, and provide mechanistic hypotheses linking alterations of promoter and enhancer activity in AF and their potential impact on atrial gene expression. Second, we obtained genome-wide DNA methylation levels at base-resolution in a subset of these samples, and identified differentially-methylated regions in AF patients that significantly associate with candidate loci. Specific transcription factor motifs are over-represented in the sequences of these hyper- and hypomethylated regions, including those corresponding to AF-associated transcription factors ETV1 [60] and HAND2 [27]. Integration of DNA methylation changes at transcription factor binding sites with H3K72ac levels and gene expression allowed us to propose co-ordinated mechanisms underlying altered gene regulation in AF for cardiovascular loci such as *LRRC4B*. Albeit with limited sample numbers and statistical power, our results represent a systematic investigation of dysregulated gene expression across the left and right atrium in persistent AF, and add to ongoing efforts to inform the genetic and epigenetic underpinnings of human population findings in AF cohorts.

Our data revealed a number of insights into how alterations in the regulatory landscape associate with persistent AF, and delineated candidate dysregulated loci and epigenomic regions. These include known AF disease loci, such as the *NPPB* locus [65, 66]. In agreement with previous work, we observed increased *NPPB* expression in AF samples, and additionally identified epigenomic regions with increased H3K27ac levels - in AF samples overall and in AF-LA; Figures 5 and S4). In contrast, other known AF disease loci such as *PITX2* did not show significant transcriptomic and epigenomic changes in our samples, which may be due to the small numbers of patients we profiled. In this regard, previous atrial gene expression datasets did not consistently observe *PITX2* expression changes in persistent AF samples [64], which have been recently validated in larger patient cohorts [23].

Among the candidate loci we identified by integration of H3K27ac and transcriptomic profiles (Figures 3 and S4), several are involved in fibrosis-induced structural remodeling, and include loci with increased (*SCX*) or decreased (*WNT5A*) activity in AF samples. SCX enables fibroblast to myofibroblast conversion, as evidenced in *in vivo* deficiency models with compromised cardiac matrix formation and loss of cardiac fibroblasts [67]. In this locus, the observed epigenomic and transcriptomic levels resulting from increased promoter activity in AF-RA samples from our cohort could be contributing to the activation of fibrotic processes in AF. Dysregulations of the Wnt signaling pathway are known to contribute to fibrotic processes in the atria, with decreased activity of most components, such as *WNT5A*, found in AF patient samples [68] and rapid-pacing in vivo models [69].

As part of our validation of candidate loci in an independent sample cohort, we qualitatively confirmed both H3K27ac enrichment and gene expression levels for loci such as *NPPB, VASH1* and *MT1X* (Figure 5). *VASH1* upregulation reportedly leads to reduced angiogenesis [70], and our findings support a model of endothelial dysfunction and a prothrombotic state promoting AF. *MT1X* decreased levels, on the other hand, are known to induce oxidative stress conditions [71], thus promoting both structural and electrical remodeling through alteration of ion channel activity.

Our DNA methylation data in these samples allowed us to identify potential mechanisms of gene dysregulation across candidate loci (Figure 6). *SLC43A3* is a membrane transporter protein responsible for uptake of purine nucleotide bases into the cells [72] as well as regulation of free fatty acids transport [73], and we found both DNA hypomethylation and chromatin hyperacetylation of its promoter in AF samples. Therefore, our findings suggest lipid metabolism dysregulation could contribute to AF disease through altered SLC43A3 expression. In the *MYLK2* gene locus, encoding a myosin light chain kinase 2 protein with crucial roles in cardiac muscle contraction [74], promoter hypomethylation in AF and increased H3K27ac levels at a putative distal enhancer contrasted with reduced *MYLK2* gene expression. Instead, our findings for this locus are consistent with the hypomethylated region regulating FOXS1 expression, which we found to be increased in AF samples. Because FOXS1 has been mechanistically implicated in liver fibrosis [75], it is tempting to speculate its upregulation in atrial tissue may contribute to fibrosis in permanent AF. Nevertheless, our observation could stem from the proposed association between AF and calcific aortic valve stenosis [76], in which FOXS1 is functionally implicated [77].

Lastly, and through investigation of overrepresented transcription factor motifs in differentially-methylated regions (Figure 7 and S7), our results provide mechanistic hypotheses by associating epigenetic dysregulation in AF with potential changes in transcription factor binding. An illustrative example is the *LRRC4B* locus, where we identified a putative *LRRC4B* enhancer hypermethylated in AF samples. In this region, one of the differentially-methylated CpGs overlaps a transcription factor motif for ETV1, a known AF-associated transcription factor [60]. This observation suggests impaired ETV1 binding in this putative enhancer could mediate reduced LRRC4B expression in AF (Figure 7). LRRC4B is a lamin-associated protein with reduced protein expression in lamin A/C haploinsufficient cardiomyocytes [61]. Our results suggest reduced *LRRC4B* expression occurs via hypermethylation of a distal enhancer [78] and reduced ETV1 binding [79, 80], potentially contributing to remodeling of atrial cardiomyocytes in AF. Nevertheless, further work would be required to substantiate this hypothesis.

There are a number of limitations to our approach. First, the sample size in our study was reduced by limited availability of atrial biopsies comprising both left and right atrial appendages from each patient. This is a common limitation in epigenomic studies on human cardiac samples, and informs our choice of semi-quantitative methods to identify hyper- and hypoacetylated epigenomic regions marked by H3K27ac [48]. Nevertheless, the integration of histone modification data with DNA methylation and transcriptomic readouts across left and right atrial samples allowed us to define candidate loci with multiple indications of gene dysregulation in AF patients – some of which we could qualitatively validate in an independent set of samples.

Second, the epigenomic profiling experiments we report targeted H3K27ac as a histone modification consistently associated with the activity of mammalian promoters and enhancers [42, 81], and DNA methylation levels as a second epigenetic modification with documented and widespread changes during AF development [38, 39]. Albeit H3K27ac is known to be dysregulated across a range of cardiovascular diseases [45, 46, 82], epigenomic regions defined by other histone modifications, such as those associated to non-canonical enhancers [83], are also likely to be modified in permanent AF. Similarly, additional epigenetic mechanisms known to be altered in AF patients, such as microRNA or long non-coding RNA expression, were not investigated in our study. Thus, our data represents a practical compromise between sample availability and epigenomic completeness. Nevertheless, denser sampling of AF patients and sinus rhythm controls would impact performance of our approach by improving sensitivity and reducing false discovery rate.

Third, our analyses used bulk readouts on cardiac tissue samples from left and right atrium, thus measuring an average of signals arising from cardiac cell types in each sample. Despite some of our identified loci containing genes with documented roles in cardiomyocytes, such as *LRRC4B*, multiple lines of evidence suggest our results are likely to inform mechanism of gene dysregulation in additional cell types. First, several of our candidate loci include genes expressed in cardiac fibroblasts and known to regulate fibrosis, such as *SCX* [84]. Second, epigenomic regions with changes in H3K27ac and DNA methylation both associate with open chromatin regions across multiple cell types in the human heart (Table S10). Although outside the scope of our work, computational deconvolution approaches [85, 86] could be applied to the datasets we report here to resolve epigenomic signals in specific cardiac cell types. The significant rise in cost notwithstanding, future studies may extend our approach by leveraging single-cell technologies for transcriptomic and epigenomic profiling [87, 88].

In summary, this study presents an integrated investigation of epigenomic and transcriptomic changes associated with persistent atrial fibrillation in the right and left atrium – thereby addressing the potential impact of dysregulated gene expression in AF susceptibility and progression. By systematically identifying epigenomic regions dysregulated in AF and their association with gene expression, we show the value of functional genomics analyses to identify candidate loci dysregulated in AF patients, and connect these changes in gene regulation with mechanistic hypotheses at the molecular level.

## MATERIALS AND METHODS

### Details of samples used in the study

Left and right adult human atrial appendages were collected from patients undergoing open-heart surgery (bypass grafting and/or atrial/mitral valve repair) at Barts Health Centre, Barts Health National Health Service (NHS) Trust. Ethical approval was obtained from the East of England-Cambridge Central Research Committee (14/EE/0007). Patients classified as SR who developed post-operative AF were excluded from the study. Samples were cryopreserved in RNAlater and stored at −80°C.

A discovery cohort of 20 samples was used for the generation of ChIP-seq and EPIC array DNA methylation data. These samples comprised 12 individuals, 7 classified as persistent AF and 5 as sinus rhythm (SR). Paired left and right atrial samples were obtained in 8 of the 12 individuals. A further 10 paired left and right atrial samples were obtained from an independent cohort of 5 individuals (3 AF and 2 SR) and used for validation of epigenome and transcriptome-wide findings in the discovery cohort through locus-level quantification of gene expression or histone modification enrichment by quantitative PCR. Details regarding patients, samples and data generated from each sample can be found in Table S1.

### Chromatin immunoprecipitation and high-throughput sequencing (ChIP-seq)

ChIP and high-throughput sequencing of the discovery cohort were performed across four different batches. The OSAT R package was used to avoid sample allocation biases and ensure even distribution of biological groups across independent batches [89]. An additional batch of ChIP was performed for the 10 samples in the validation cohort for downstream qPCR experiments. All ChIP experiments were performed as described in existing protocols with minor modifications [90], using commercial antibodies for H3K27ac (ab4729, Abcam) and H3K4me3 (05-1339, Sigma-Aldrich). The latter antibody was only used for a subset of samples (as detailed in our GEO dataset, see Data Availability section).

### Computational analysis of ChIP-seq data

#### Basic alignment and peak calling

Aligned bam files were obtained with bwa 0.7.17 and the Ensembl v114 hg38 human genome assembly. We used macs2 [91] to call peaks for each ChIP-Seq replicate, using default parameters and “--keep-dup all” to retain duplicate reads. Before peak-calling, multi-mapping reads were removed and read-depth adjusted to 20 million uniquely mapped reads (or all available reads for low-depth libraries).

#### Definition of regulatory regions across each sample type

we first constructed sets of reproducible peaks for each of the four sample groups by calling peaks with Genrich (https://github.com/jsh58/Genrich) on all ChIP and input libraries within each group. We then used the chip-greylist tool to identify peaks reproducibly called from input samples alone (“greylisted”), which we excluded from analysis.

### Definition of enriched H3K27ac regions across sample groups from pairwise replicate comparisons

H3K27ac-enriched peaks identified with MACS2 within individual samples were used to define a consensus set of enriched regions in each of the four biological categories (AF-LA, AF-RA, SR-LA and SR-RA) or the two overall disease groups (AF and SR). To do this, regions with differential signal were identified for pairwise comparisons across all individual samples in the dataset (for n = 20, 190 possible comparisons) using MACS2 ‘bdgdiff’ function with a likelihood ratio cutoff of 100 (c = 2), 300bp of differential region minimum size (l = 300) and 100 bp maximum gap between peaks to be merged into a single one (g = 100). For each pairwise comparison, two peaksets were obtained (corresponding to peaks enriched in either sample pair, 190×2 = 380 enriched peaksets). These peaksets were then used to build a consensus set of enriched regions for each category (excluding within-category pairwise comparisons). Enriched peaks within each category with at least 1bp overlap were merged into a single consensus region. To achieve similar numbers of enriched regions across sample groups, the number of sets required to overlap for a region to be included in the consensus set was adjusted relative to the number of replicates within each category. To exclude enriched regions resulting from merging and concatenation, at least 30% of the length of enriched peaks was required to reference peaks collectively called for all replicates in each category (obtained as detailed above with Genrich). Lastly, enriched regions overlapping multiple categories were removed, retaining uniquely enriched regions for further analyses.

To obtain normalised coverage values, we used the “cov” function in BAMscale [92] to calculate FPKM read coverage in H3K27ac ChIP and input samples for enriched regions across sample groups, as described previously [44]. H3K27ac coverage was then normalised by input coverage and quantile normalisation, and represented in logarithmic scale.

### RNA-sequencing analyses of differential gene expression

Raw RNA-seq data generated for a former study [33] in largely overlapping samples (Table S1) was obtained from the National Centre for Biotechnology Information’s Gene Expression Omnibus (GEO) (GSE128188) and re-analysed to identify DEGs between sample groups.

Reads were aligned to the hg38 reference human genome using STAR with default parameters [93]. A count matrix of all reads was created with featureCounts [94] and differential expression analysis was performed using DESeq2 (v1.42.1) [95]. After incorporating Ensembl v114 bioMart human gene annotations [96], protein-coding genes with at least 5 read counts across all samples were selected for downstream analysis (n = 18,370 genes). Read counts were normalised by library size factors, and p-values were calculated using a negative bimodal distribution and corrected for multiple testing using Benjamini and Hochberg method.

Integration of DEGs with ChIP-seq H3K27ac enrichment data was investigated by exploring the overlap of DEGs and genes in proximity to H3K27ac-enriched regions using the default gene-to-peak association rule in GREAT (v4.0.4) [97]. Significance was tested with Fisher’s Exact Test, with all genes used in differential expression analysis as background.

### Experimental validation of selected candidate regions in an independent patient cohort

Validation of gene expression and H3K27ac enrichment levels across candidate loci in an independent patient cohort was performed with qPCR technology. For RT-qPCR validation, gene-specific primers were designed with NCBI Primer Designing tool (Table S9). Total RNA was isolated from 30-80 mg of atrial tissue. Complete disruption and homogenisation of tissue was achieved with 3 cycles of 15 seconds at 4.5 m/s in a FastPrep-24 bead beating and lysis system (MP Biomedicals). The RNeasy Plus Universal kit (Qiagen) and TRIzol reagent (Ambion) were used for RNA extraction. RNA integrity was assessed with the RNA 6000 Nano Kit for Bioanalyzer 2100 (Agilent). 1000 ng of RNA were reversed transcribed into complementary DNA (cDNA) using the High-Capacity cDNA Reverse Transcription Kit (Fisher Scientific), and RT-qPCR reactions were performed using 5 ng of template cDNA. Ct values for each gene of interest were normalised to those of *TPB* reference gene using the ΔCt method [98].

H3K27ac ChIP of atrial samples in the validation cohort was performed as above. Qpcr primers were designed to target selected enriched regions identified in the discovery cohort’s ChIP-seq data, with the NCBI Primer Designing tool (Table S9). ChIP and input DNA samples were diluted 1:5 and 1:10 respectively for qPCR reactions. ChIP enrichments were normalised to those of a region in the *GAPDH* reference locus over input samples (2^ - (Ct_Target_ – Ct*_GAPDH_*)_ChIP_ – (Ct_Target_ – Ct*_GAPDH_*)_Input_).

Universal KAPA SYBR reagents (KAPA Biosystems) were used for all qPCR reactions on a Roche LightCycler480 system, with the following parameters: 95°C for three minutes (pre-incubation), 40 cycles at 95°C for 10 seconds, 60°C for 20 seconds and 72°C for 1 second. Generation of melt curves after 40 cycles was carried out to control for amplification of non-specific products.

### Experimental assessment of DNA methylation levels with EPIC array technology

Genomic DNA was isolated from 16 atrial samples (AF-LA, n=6; AF-RA, n=5; SR-LA, n=4; SR-RA, n=4) using the DNeasy Blood & Tissue Kit (Qiagen). Bisulfite treatment and subsequent amplification and hybridisation to DNA methylation profiling microarray was performed by the QMUL Genome Centre. The Infinium MethylationEPIC v2.0 kit was used, providing coverage of approximately 930K CpGs, including RefSeq annotated genes, CpG islands and enhancer regions. Even and random distribution of biological groups across two chips of 8 samples each was allocated using the OSAT R Bioconductor package [89].

### Computational analysis of DNA methylation data

We quantified methylation levels for a total of 930,698 CpG sites using the R Bioconductor package *meffil* (v1.3.6) [99]. Briefly, functional normalisation was applied to array intensity data (“.idat” files) to correct for between-array technical variation. Low-quality and sex chromosome probes were removed. Methylation status was reported for all measured CpG sites as β-values ranging from 0 to 1 representing the fraction of DNAm signal compared to the total locus signal intensity.

An EWAS model was constructed to test for association of DNA methylation CpG sites with disease status, incorporating surrogate variables to correct for confounders. Details about the genomic locations of all CpG sites were loaded from the Illumina methylation EPIC array v2.0 annotation package (IlluminaHumanMethylationEPICv2anno.20a1.hg38). Differentially methylated regions (DMRs) between AF and SR samples were called with the *dmrff* package (https://github.com/perishky/dmrff) by combining the EWAS summary statistics from CpG sites located close together and showing strong associations. DMRs were defined as regions containing more than one CpG with at most 500 bp between consecutive sites and effect estimates with the same direction. Significant DMRs were called with p-values < 0.05, with Bonferroni multiple-testing correction.

### Analyses of enriched transcription factor motifs, gene ontology terms and binding regions

For *de novo* and known motif enrichment analysis, the findMotifsGenome.pl script (-size given) from HOMER v5.1 [54] was used on H3K27ac-enriched sets of interest and hyper- or hypomethylated DMRs, using the sets of reference peaks from each category and all candidate regions with aggregated CpG methylation signal respectively as background. In addition, significance for TF binding sites from ChIP-seq experimental data generated from the ReMap catalogue was assessed using the shuffling procedure implemented in the ReMapEnrich R package [55]. A subset of ChIP-seq TF binding experiments from the ReMap catalogue was selected from biologically relevant samples (i.e., human cardiac tissue or *in vitro* heart models).

For enrichment of AF-associated TFs on the DMRs, these were selected from recent GWAS [19] and binding motifs were collected from JASPAR [100]. FIMO online tool (Find Individual Motif Occurrences) from the MEME suite (v.5.5.7) (match p-value < 0.001) was used to annotate motifs over DMRs and the background set of regions [101]. A hypergeometric test was used to calculate statistical significance of motif occurrences across hypermethylated and hypomethylated regions (Bonferroni correction was applied for multiple testing).

For gene ontology enrichment analyses, genes were assigned to enriched regions based on the default association rule in GREAT v4.0.4 [102], and evaluated for ontology term enrichments against reference peaks from each category using the enrichGO function from clusterProfiler v4.15.1 [103].

### Enrichment of GWAS signals and cell-type specific chromatin accessibility

Genetic lead variants associated to cardiovascular traits at genome-wide significance (p-value < 10^-8^) were obtained from the NHGRI-EBI GWAS catalog [56] (version e113, 2025), for all heart-related traits (search term “heart” across associated traits in the catalog), and four major cardiovascular diseases and traits (search terms “atrial fibrillation, “QT interval”, “coronary artery disease” and “cardiomyopathy”). Each set of lead variants was annotated with proxy variants in strong LD (R^2^ > 0.8) using SNIPA [104]. The enrichment of GWAS SNPs in heart promoters and enhancers across epigenomically-conserved, primate-specific and human only categories calculated with a hypergeometric test.

Signatures of gene regulation across heart cell types and cardiomyocytes were obtained from previously published datasets: RPKM chromatin accessibility levels across cardiac cell types [58], and significant promoter-enhancer interactions in human iPSCs and iPSC-derived cardiomyocytes were obtained from [57].

## Supporting information

Supplemental Data document incl. figures and two tables

## AUTHOR CONTRIBUTIONS

AR, SF, DFA, JC, GT and DV designed and performed experiments. AR, CGB and DV analysed data. AMT, AT and PBM provided essential reagents and technical advice. DV secured funding and coordinated the study. DV and AR wrote the manuscript. All authors read and approved the manuscript.

## ACKNOWLEDGEMENTS

We thank the QMUL Genome Centre for assistance with DNA methylation analyses on EPIC v2 arrays and members of the QMUL Centre for Epigenetics, Prof. Miguel Manzanares (CBMSO, Madrid, Spain) and Dr. Ines Cebola (Imperial College London, UK) for helpful criticism and discussions.

## DECLARATIONS

### ETHICAL APPROVAL AND CONSENT TO PARTICIPATE

Research involving human participants was conducted in accordance with the Declaration of Helsinki. Ethical approval was obtained in collaboration with the National Institutes for Health Research (NIHR) Barts Biomedical Research Centre Cardiovascular Bio-Registry from the East of England-Cambridge Central Research Committee (14/EE/0007). Patients at Barts Heart Centre, Barts Health National Health Service (NHS) Trust gave informed consent to participate in the study.

### AVAILABILITY OF DATA AND MATERIALS

The ChIP-sequencing and DNA methylation data reported here has been deposited to GEO with accession numbers GSE227793 and GSE291249, respectively.

Analysis code is available in Github: https://github.com/adrian-rodr/Rodriguez_2025_AFepigenome.git

