## Supplemental Data document incl. figures and two tables for "An epigenomic investigation of atrial fibrillation in a matched left and right atrial human cohort"

**SUPPLEMENTAL INFORMATION**

**Integrated epigenomic analysis in a paired left and right atrial fibrillation cohort**
Rodriguez et al. 2025

---

#### SUPPLEMENTAL DATA


**Figure S1. Experimental design summary and extended details on H3K27ac profiling** (related to Figure 1) **(page 3)**

**Figure S2. Methodological details and extended properties for H3K27ac epigenomic regions enriched across sample groups** (related to Figure 2) **(p. 5)**

**Figure S3. Additional analyses on H3K27ac-enriched regions across sample groups (overlap with transcription factor motifs, binding events and cardiovascular GWAS variants)** (related to Figure 2) **(p. 7)**

**Figure S4. Gene expression re-analysis and extended details on candidate loci identified from integration of epigenomic and transcriptomic signals** (related to Figure 3) **(p. 9)**

**Figure S5. Extended details and analyses of candidate loci in an independent replication cohort** (related to Figures 4 and 5) **(p. 11)**

**Figure S6 EWAS results and additional properties of DMRs across SR and AF samples** (related to Figure 6) **(p. 13)**

**Figure S7. Additional details and analyses for DMRs and their overlap with transcription factor motifs, binding events and GWAS variants** (related to Figure 7) **(p. 15)**

**Table S1**: Sample and patient characteristics, including experimental readouts obtained in each sample. **(p. 17)**

**Table S2.** Differentially-enriched regions with standard methods (DiffBind, TCseq). [SupplementalTable_S2.xlsx]

**Table S3**. Genomic regions enriched in H3K27ac across the four sample groups, and in AF vs SR samples. [SupplementalTable_S3.xlsx]

**Table S4.** Differentially-expressed genes across anatomical sides, disease status, or in each of the four sample groups. [SupplementalTable_S4.xlsx]

**Table S5**. Candidate loci with hallmarks of altered gene regulation in atrial fibrillation patients. [SupplementalTable_S5.xlsx]

**Table S6**. Candidate loci selected for validation in an independent replication cohort with locus-specific quantitative PCR. **(p. 18)**

**Table S7**. Single CpG and DMR regions showing DNA methylation differences between AF and SR samples. [SupplementalTable_S7.xlsx]

**Table S8**. Subset of candidate loci with proximal DMRs. [SupplementalTable_S8.xlsx]

**Table S9**. RT-QPCR and ChIP-QPCR primers used for targeted validation of candidate regions. [SupplementalTable_S9.xlsx]

**Table S10**. Overlap of chromatin accessibility regions across cardiac cell types with enriched H3K27ac regions and DMRs. [SupplementalTable_S10.xls]

#### SUPPLEMENTAL FIGURES

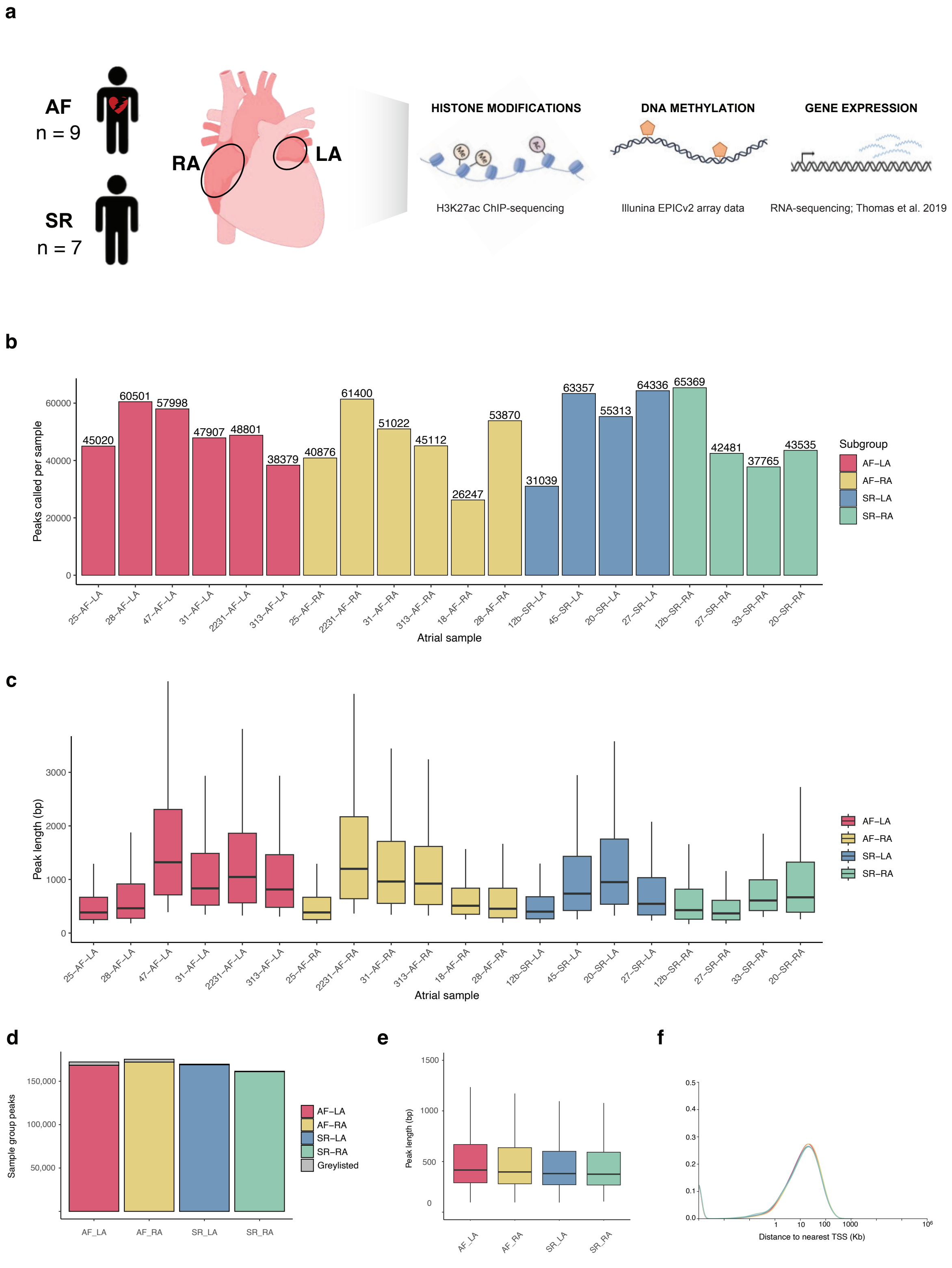


**Figure S1. Experimental design summary and extended details on H3K27ac profiling** (related to Figure 1)

**a.** Summary of experimental design in the study. Atrial fibrillation (AF) or sinus rhythm (SR) atrial samples from the right (RA) and left atrium (LA) were primarily used for H3K27ac ChIP-sequencing to profile active promoters and enhancers across individuals and sample groups. Most of these samples had been previously used for transcriptome profiling, and we additionally obtained DNA methylation EPIC array data for a subset of samples. See Table S1 for full details.

**b.** Numbers of H3K27ac peaks called in each sample (Methods). Samples are coloured by sample group, with a similar average number of peaks obtained for each (AF-LA: 49,768; AF-RA: 46,421; SR-LA: 53,511; SR-RA: 47,288).

**c.** Peak lengths for each individual sample are represented as box plots, and coloured according to sample group. Average lengths per sample group: AF-LA, 1,189 bp; AF-RA, 1,111; SR-LA, 1,024; SR-RA, 740.

**d.** Barplot with number of peaks consistently called in samples from each group, as obtained with Genrich (“reference peak set”, Methods). Peaks called in one or more input samples from each group were excluded from this set (“greylisted”, grey bars).

**e**. Lengths for reference peaks in each sample group. Average lengths: AF-LA, 991 bp; AF-RA, 905; SR-LA, 763; SR-RA, 669.

**f**. Density plots showing distance to annotated transcriptional start sites (TSSs) for reference peaks in each sample group. As expected, most H3K27ac reference peaks are distal to annotated TSSs.

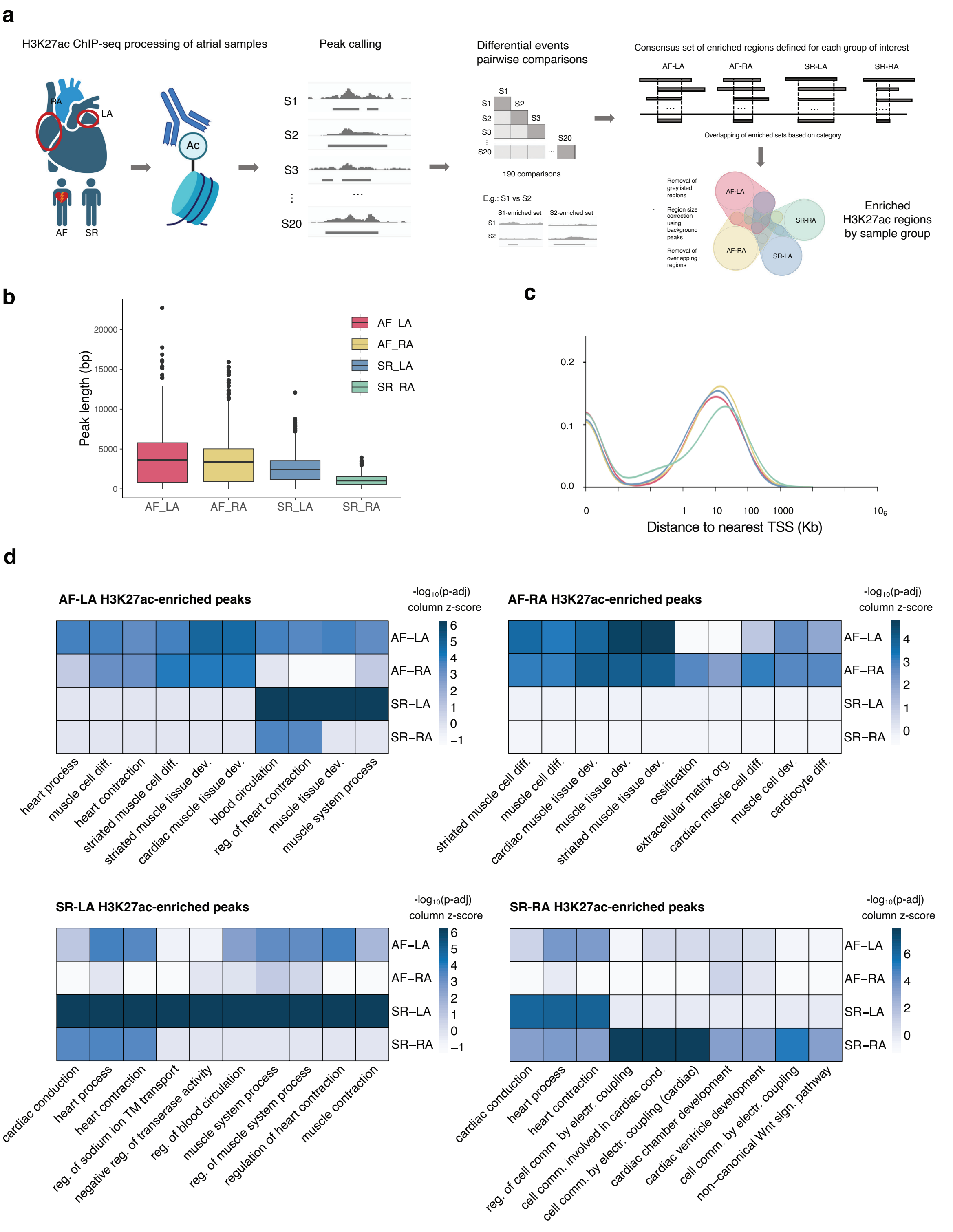


**Figure S2. Methodological details and extended properties for H3K27ac epigenomic regions enriched across sample groups** (related to Figure 2)

**a.** Summary diagram of the approach used to identify enriched H3K27ac genomic regions in each sample group. Peak calling: H3K27ac ChIP-sequencing data was processed to call regions of enrichment (peaks) in each individual sample. Differential pairwise comparisons:

quantitative H3K27ac levels were then compared across each pair of samples to identify sets of regions enriched in either of the two samples for each pair. Consensus sets of enriched regions: a set of enriched H3K27ac regions was then defined in each sample group by merging enriched pairwise sets, requiring each consensus region to be present in at least 15-50% of comparisons (Methods). These sets of enriched regions were then compared, with those overlapping more than one sample group excluded from analysis.
**b**. Boxplot representation for lengths of enriched H3K27ac regions in each sample group. Average lengths: AF-LA, 3,972; AF-RA, 3,488; SR-LA, 2,517; SR-RA, 1,103.
**c**. Distributions of distance to the nearest annotated transcriptional start site (TSS) for enriched H3K27ac regions in each sample group
**d**. Heatmap of gene ontology terms (Biological process) associated with enriched H3K27ac regions in each sample group. P-adjusted values indicate significance of GO category enrichment calculated using a hypergeometric test after FDR correction (enrichGO function from clusterProfiler).
See also Tables S2 and S3.

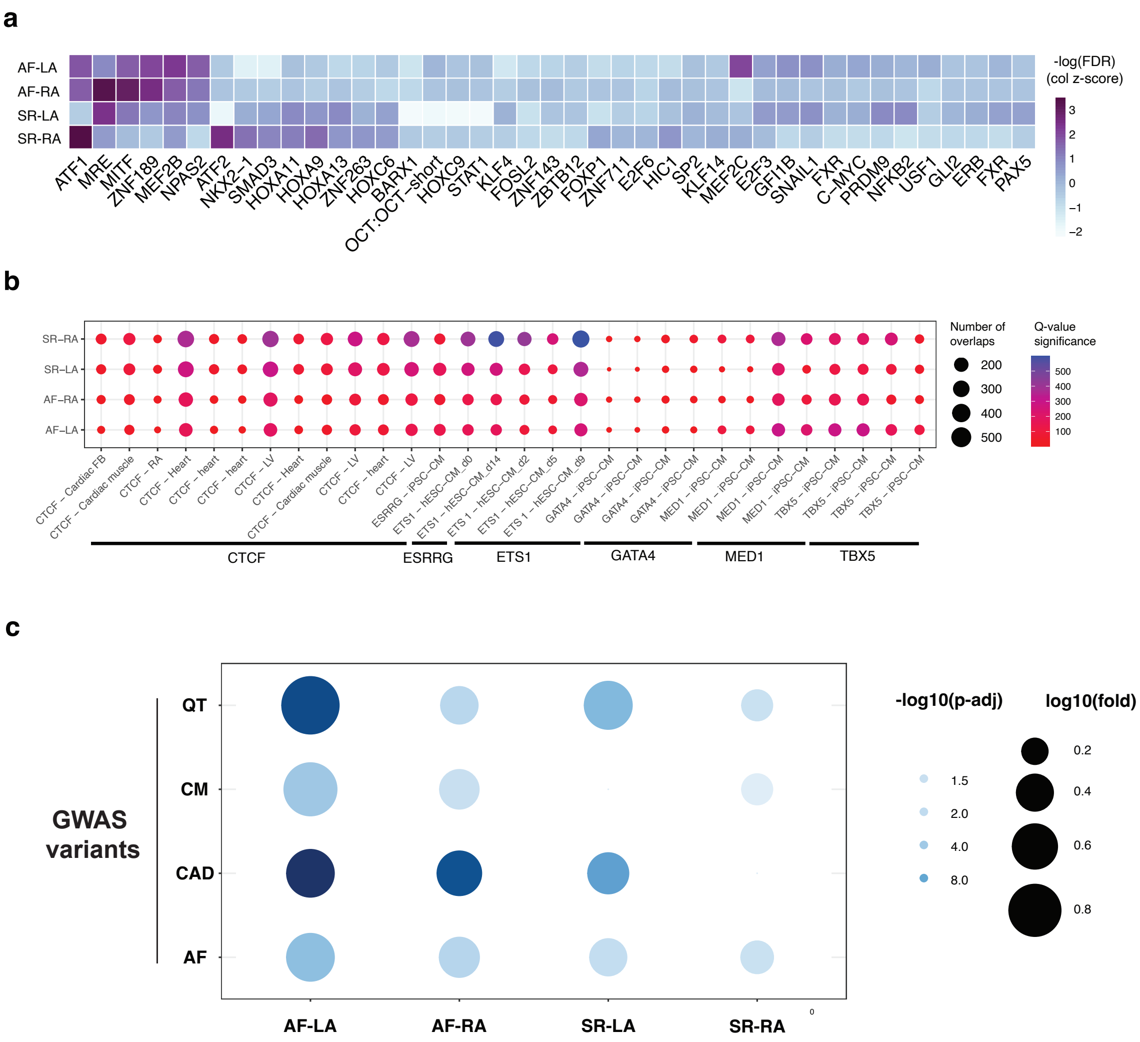


**Figure S3. Additional analyses on H3K27ac-enriched regions across sample groups (overlap with transcription factor motifs, binding events and cardiovascular GWAS variants)** (related to Figure 2)

**a**. Heatmap of transcription factor binding motifs (TFBSs) overrepresented in the DNA sequences of enriched H3K27ac regions for each sample group. Significance levels are illustrated in -log10(FDR) scale. These were estimated based on resemblance with known motif databases using the HOMER enrichment test assuming binomial distribution of motif occurrences.

**b.** Selected transcription factor binding events overrepresented in enriched H3K27ac regions for each sample group are represented as a circle plot. Binding data corresponds to ChIP-sequencing sets reported in cardiac tissue or stem-cell derived cardiomyocytes (iPSC-CM or hESC-CM), as catalogued in Remap 2022. Circle sizes denote the number of overlaps, and colour scale the Q-value of significance (FDR-corrected p-value from Fisher's Exact Test).

**c.** Cardiovascular GWAS variants over-represented in H3K27ac-enriched regions. Circle plot represents fold enrichment (circle size, log10 fold) and statistical significance (circle shade, -log10 of adjusted p-values, hypergeometric test with BH correction for multiple comparisons) for the enrichment of cardiovascular GWAS variants in H3K27ac-enriched regions across sample groups (AF-LA, AF-RA, SR-LA and SR-RA, x-axis). GWAS variants correspond to all lead variants in the NIHR-EBI GWAS catalog plus proxy variants in strong LD for atrial fibrillation (AF), coronary artery disease (CAD), cardiomyopathies (CM) and QT interval duration (QT).

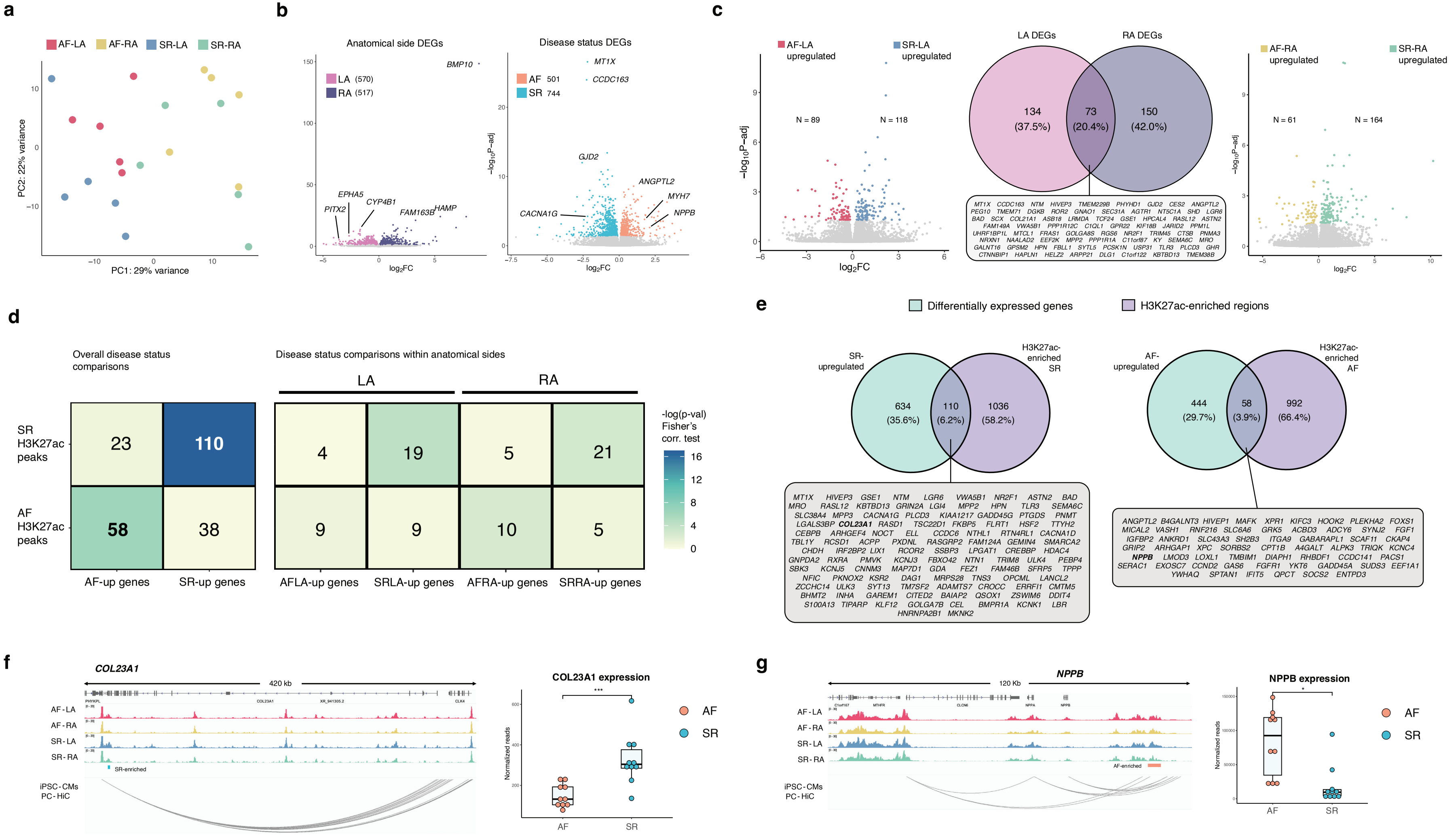
a
AF-LA
AF-RA
SR-LA
SR-RA
PC1: 29% variance
PC2: 22% variance
b
Anatomical side DEGs
LA (570)
RA (517)
BMP10
EPHA5
PITX2
CYP4B1
FAM163B
HAMP
log2FC
Disease status DEGs
AF 501
SR 744
MT1X
CCDC163
GJD2
ANGPTL2
MYH7
NPPB
CACNA1G
−log10P−adj
c
AF-LA upregulated
SR-LA upregulated
N = 89
N = 118
LA DEGs
RA DEGs
134 (37.5%)
73 (20.4%)
150 (42.0%)
MT1X CCDC163 NTM HIVEP3 TMEM229B PHYHD1 GJD2 CES2 ANGPTL2 PEG10 TMEM71 DGKB ROR2 GNAO1 SEC31A AGTR1 NT5C1A SHD LGR6 BAD SCX COL21A1 ASB18 LRMDA TCF24 GSE1 HPCAL4 RASL12 ASTN2 FAM149A VWA5B1 PPP1R12C C1QL1 GPR22 KIF18B JARID2 PPM1L UHRF1BP1L MTCL1 FRAS1 GOLGA8S RGS6 NR2F1 TRIM45 CTSB PNMA3 NRXN1 NAALAD2 EEF2K MPP2 PPP1R1A C11orf87 KY SEMA6C MRO GALNT16 GPSM2 HPN FBLL1 SYTL5 PCSK1N USP31 TLR3 PLCD3 GHR CTNNBIP1 HAPLN1 HELZ2 ARPP21 DLG1 C1orf122 KBTBD13 TMEM38B
AF-RA upregulated
SR-RA upregulated
N = 61
N = 164
d
Overall disease status comparisons
Disease status comparisons within anatomical sides
LA
RA
SR H3K27ac peaks
AF H3K27ac peaks
23
110
58
38
4
19
5
21
9
9
10
5
AF-up genes
SR-up genes
AFLA-up genes
SRLA-up genes
AFRA-up genes
SRRA-up genes
-log(p-val) Fisher's corr. test
16
12
8
4
0
e
Differentially expressed genes
H3K27ac-enriched regions
SR-upregulated
H3K27ac-enriched SR
634 (35.6%)
110 (6.2%)
1036 (58.2%)
MT1X HIVEP3 GSE1 NTM LGR6 VWA5B1 NR2F1 ASTN2 BAD MRO RASL12 KBTBD13 GRIN2A LGI4 MPP2 HPN TLR3 SEMA6C SLC38A4 MPP3 CACNA1G PLCD3 KIAA1217 GADD45G PTGDS PNMT LGALS3BP COL23A1 RASD1 TSC22D1 FKBP5 FLRT1 HSF2 TTYH2 CEBPB ARHGEF4 NOCT ELL CCDC6 NTHL1 RTN4RL1 CACNA1D TBL1Y RCSD1 ACPP PXDNL RASGRP2 FAM124A GEMIN4 SMARCA2 CHDH IRF2BP2 LIX1 RCOR2 SSBP3 LPGAT1 CREBBP HDAC4 GNPDA2 RXRA PMVK KCNJ3 FBXO42 NTN1 TRIM8 ULK4 PEBP4 SBK3 KCNJ5 CNNM3 MAP7D1 GDA FEZ1 FAM46B SFRP5 TPPP NFIC PKNOX2 KSR2 DAG1 MRPS28 TNS3 OPCML LANCL2 ZCCHC14 ULK3 SYT13 TM7SF2 ADAMTS7 CROCC ERRFI1 CMTM5 BHMT2 INHA GAREM1 CITED2 BAIAP2 QSOX1 ZSWIM6 DDIT4 S100A13 TIPARP KLF12 GOLGA7B CEL BMPR1A KCNK1 LBR HNRNPA2B1 MKNK2
AF-upregulated
H3K27ac-enriched AF
444 (29.7%)
58 (3.9%)
992 (66.4%)
ANGPTL2 B4GALNT3 HIVEP1 MAFK XPR1 KIFC3 HOOK2 PLEKHA2 FOXS1 MICAL2 VASH1 RNF216 SLC6A6 GRK5 ACBD3 ADCY6 SYNJ2 FGF1 IGFBP2 ANKRD1 SLC43A3 SH2B3 ITGA9 GABARAPL1 SCAF11 CKAP4 GRIP2 ARHGAP1 XPC SORBS2 CPT1B A4GALT ALPK3 TRIQK KCNC4 NPPB LMOD3 LOXL1 TMBIM1 DIAPH1 RHBDF1 CCDC141 PACS1 SERAC1 EXOSC7 CCND2 GAS6 FGFR1 YKT6 GADD45A SUDS3 EEF1A1 YWHAQ SPTAN1 IFIT5 QPCT SOCS2 ENTPD3
f
COL23A1
420 kb
AF-LA
AF-RA
SR-LA
SR-RA
SR-enriched
iPSC-CMs PC-HiC
COL23A1 expression
***
Normalized reads
AF
SR
g
NPPB
120 Kb
AF-enriched
NPPB expression
*

**Figure S4. Gene expression re-analysis and extended details on candidate loci identified from integration of epigenomic and transcriptomic signals** (related to Figure 3)

**a.** Principal component analysis of gene expression levels across individual samples in AF-LA, AF-RA, SR-LA and SR-RA groups, showing the first two principal components as a scatter plot. Variance across PC1 and PC2 provides a degree of separation across anatomical side (LA samples on the left and RA samples on the right).
**b.** Volcano plots of differentially expressed genes (DEGs) in each anatomical side (LA, 570 upregulated genes; RA, 517 upregulated genes) and disease status (AF, 501 upregulated genes; SR, 744 upregulated genes).
**c.** Volcano plots of differentially expressed genes across disease status, either in left atrium (left plot: AF-LA upregulated, 89 genes; AF-RA upregulated, 118 genes) or right atrium samples (right plot: AF-RA upregulated, 61 genes; SR-RA upregulated, 164 genes). The central Venn diagram corresponds to overlap of genes found to be differentially expressed in each anatomical side.
**d.** Association of H3K27ac enriched regions in each sample group with differentially expressed genes, either for AF vs SR samples (left plot) or for enriched regions / DEGs across all four sample groups (right plot). Numbers in each cell represent overlapping gene loci, and colour shades statistical significance ($-\log_{10}$(p-value), Fisher's exact test).
**e.** Definition of candidate loci with altered gene regulation in AF or SR samples. Venn diagrams represent the overlap between upregulated genes (cyan) and enriched H3K27ac regions (violet), either in SR (left) or AF (right). Grey insets under each Venn diagram detail the candidate loci derived from each comparison.
**f.** and **g.** Examples of candidate loci from e. **f.** (Left) H3K27ac average fold enrichment over input for each sample group across the *COL23A1* locus, including an SR-enriched H3K27ac region (left, SR-enriched). Promoter-capture HiC data in iPSC-cardiomyocytes supports 3D interactions between this region and a distal region downstream. (Right) *COL23A1* expression levels are upregulated in SR samples, in concordance with the H3K27ac enriched region (***, $p < 0.001$; DESeq2 adjusted p-values) **g**. (Left) Representation as in f. across the *NPPB* locus, including an AF-enriched H3K27ac region with supporting evidence of chromatin contacts with promoters in this locus. (Right) *NPPB* expression levels are significantly higher in AF samples (*, $p < 0.05$; DESeq2 adjusted p-values).
See also Tables S4 and S5.

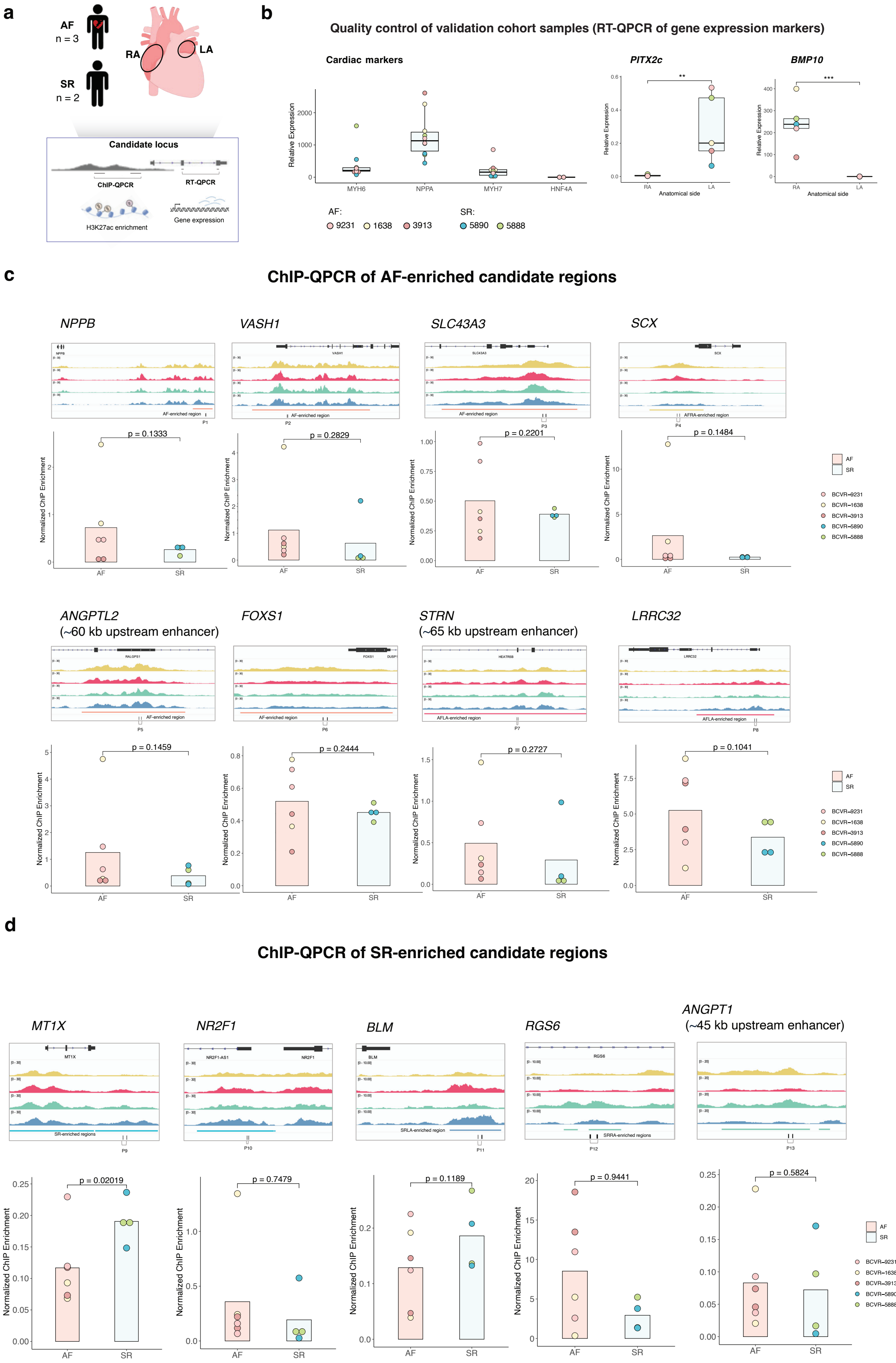
a
AF
n = 3
SR
n = 2
RA
LA
Candidate locus
ChIP-QPCR
RT-QPCR
H3K27ac enrichment
Gene expression
b
Quality control of validation cohort samples (RT-QPCR of gene expression markers)
Cardiac markers
PITX2c
BMP10
Relative Expression
MYH6
NPPA
MYH7
HNF4A
Anatomical side
AF:
9231
1638
3913
SR:
5890
5888
c
ChIP-QPCR of AF-enriched candidate regions
NPPB
VASH1
SLC43A3
SCX
p = 0.1333
p = 0.2829
p = 0.2201
p = 0.1484
Normalized ChIP Enrichment
AF
SR
BCVR–9231
BCVR–1638
BCVR–3913
BCVR–5890
BCVR–5888
ANGPTL2
(~60 kb upstream enhancer)
FOXS1
STRN
(~65 kb upstream enhancer)
LRRC32
p = 0.1459
p = 0.2444
p = 0.2727
p = 0.1041
d
ChIP-QPCR of SR-enriched candidate regions
MT1X
NR2F1
BLM
RGS6
ANGPT1
(~45 kb upstream enhancer)
p = 0.02019
p = 0.7479
p = 0.1189
p = 0.9441
p = 0.5824

**Figure S5. Extended details and analyses of candidate loci in an independent replication cohort** (related to Figures 4 and 5)

**a.** Summary diagram of the experimental approach used for independent validation of candidate loci. A set of 3 AF and 2 SR samples comprising both left and right atrium were used for locus-specific quantitative PCR validation of gene expression levels (RT-QPCR) or H3K27ac enrichment (ChIP-QPCR) (Methods).

**b.** Quality control of independent validation samples by RT-QPCR. Expression levels were assessed for cardiac markers (left: *MYH6*, *NPPA* and *MYH7*, using liver-specific *HNF4A* as a control), a left atrium marker (*PITX2*) and a right atrium marker (*BMP10*). P-values: ***, $p < 0.001$; ** $p < 0.01$ (Student's t-test)

**c.** ChIP-QPCR validation of H3K27ac enrichment across selected AF-enriched candidate regions. For each region, top genome tracks show average H3K27ac fold enrichment over input in each sample group in the discovery cohort, enriched candidate regions (coloured bars, bottom), and the location of QPCR primers (e.g. P1). Bottom barplots show normalised H3K27ac ChIP enrichment (Methods) for each assessed region across validation samples. P-values correspond to one-sided t-test.

**d.** Representation and details as in c. for ChIP-QPCR validation of H3K27ac enrichment across selected SR-enriched candidate regions. P-values correspond to one-sided t-test. See also Tables S6 and S9.

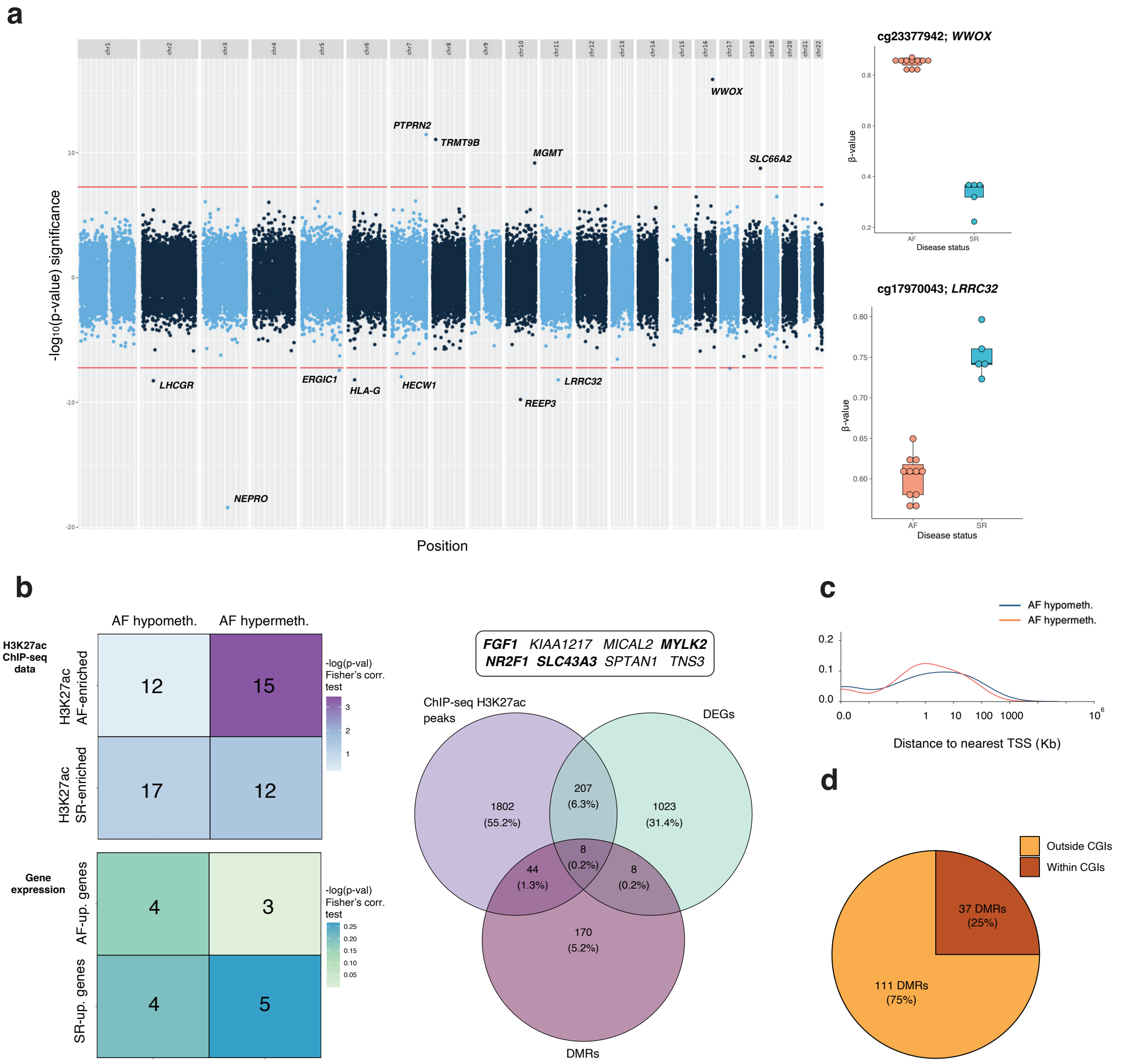


**Figure S6. EWAS results and additional properties of DMRs across SR and AF samples** (related to Figure 6)

**a.** Manhattan plot for EWAS analysis of EPIC v2 array data across SR and AF samples. Dashed red lines indicate the statistical threshold for genome-wide significance ($p < 10^{-8}$), and labelled points above and below this line correspond to CpGs with methylation levels significantly higher in AF (above) or SR samples (below). Boxplots on the right provide two examples of genome-wide differentially methylated CpGs (DMCs) for the *WWOX* and *LRRC32* loci (respectively).

**b.** Association of differentially-methylated regions (DMRs) with H3K27ac enriched regions and differentially expressed genes in AF and SR samples. (Left) Heatmaps show the number of gene loci associated to both DMRs and H3K27ac enriched regions (top), or DMRs and differentially expressed genes (bottom). Colour scales correspond to statistical significance ($-\log_{10}$(p-value), Fisher's exact test). (Right) Venn diagram showing the overlap of gene loci associated to enriched H3K27ac regions, differentially-expressed genes (DEGs) and DMRs (p=0; Chi-square test of independence). In all cases, gene loci correspond to genes associated to differential regions between AF and SR samples (e.g. both AF-upregulated and SR-upregulated genes).

**c.** Distribution of distances to the nearest annotated TSS for DMRs hypermethylated (red) or hypomethylated (blue) in AF samples.

**d.** Annotation of DMRs according to CpG island annotations (CGIs; Methods), as either within (brown) or outside (orange) CGIs.

See also Tables S7 and S8.

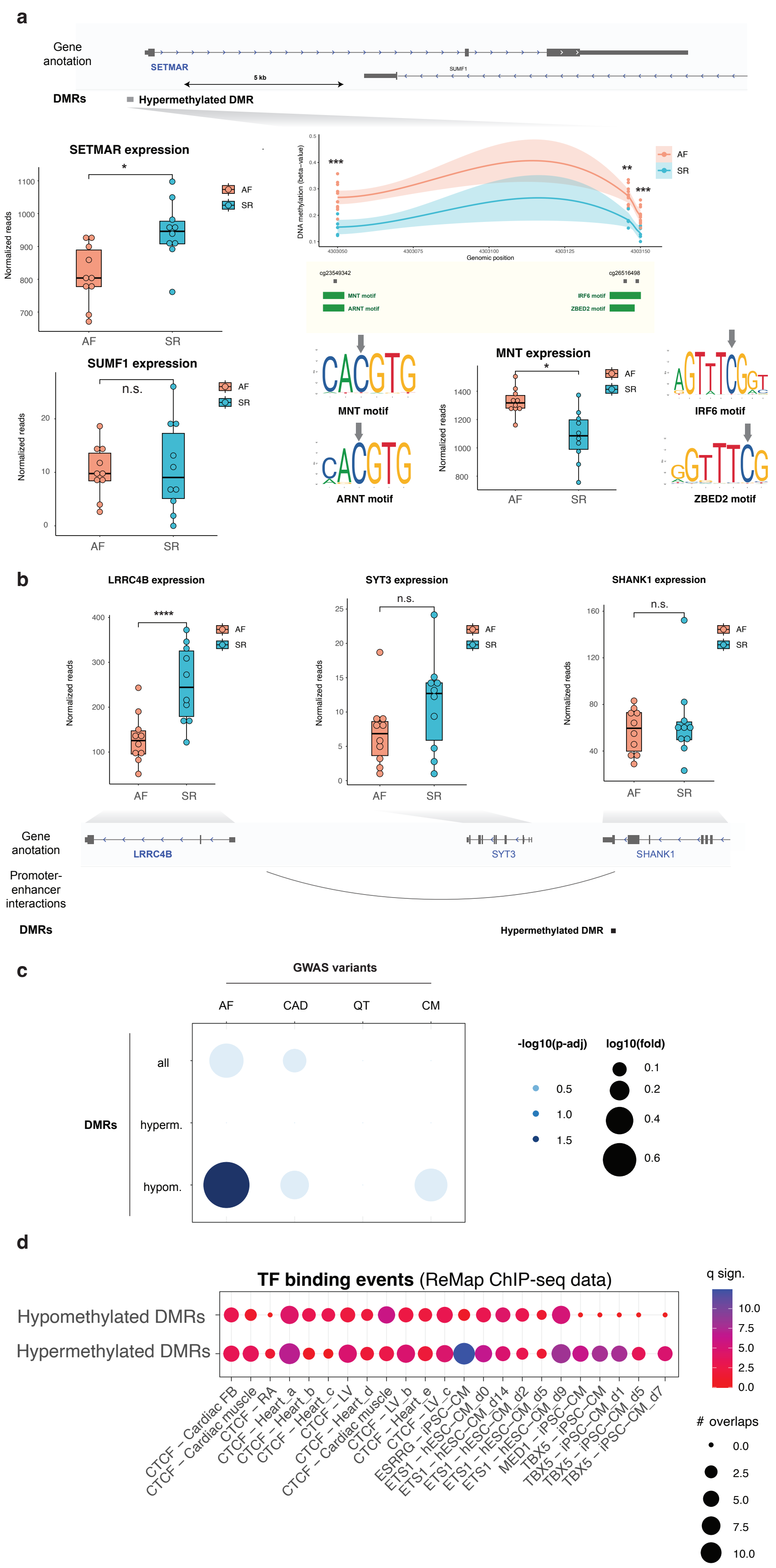
a
Gene anotation
SETMAR
SUMF1
5 kb
DMRs
Hypermethylated DMR
SETMAR expression
Normalized reads
AF
SR
*
SUMF1 expression
n.s.
DNA methylation (beta-value)
Genomic position
***
**
***
cg23549342
cg26516498
MNT motif
ARNT motif
IRF6 motif
ZBED2 motif
MNT expression
b
LRRC4B expression
****
SYT3 expression
n.s.
SHANK1 expression
n.s.
Gene anotation
LRRC4B
SYT3
SHANK1
Promoter-enhancer interactions
DMRs
Hypermethylated DMR
c
GWAS variants
AF
CAD
QT
CM
all
hyperm.
hypom.
DMRs
-log10(p-adj)
0.5
1.0
1.5
log10(fold)
0.1
0.2
0.4
0.6
d
TF binding events (ReMap ChIP-seq data)
Hypomethylated DMRs
Hypermethylated DMRs
q sign.
10.0
7.5
5.0
2.5
0.0
### overlaps
0.0
2.5
5.0
7.5
10.0
CTCF – Cardiac FB
CTCF – Cardiac muscle
CTCF – RA
CTCF – Heart_a
CTCF – Heart_b
CTCF – Heart_c
CTCF – LV
CTCF – Heart_d
CTCF – Cardiac muscle
CTCF – LV_b
CTCF – Heart_e
CTCF – LV_c
ESRRG – iPSC–CM
ETS1 – hESC–CM_d0
ETS1 – hESC–CM_d14
ETS1 – hESC–CM_d2
ETS1 – hESC–CM_d5
ETS1 – hESC–CM_d9
MED1 – iPSC–CM
TBX5 – iPSC–CM
TBX5 – iPSC–CM_d1
TBX5 – iPSC–CM_d5
TBX5 – iPSC–CM_d7

**Figure S7. Additional details and analyses for DMRs and their overlap with transcription factor motifs, binding events and GWAS variants** (related to Figure 7)

**a.** Example of a hypermethylated DMR in the SETMAR locus (DMR49), for differentially-methylated CpGs overlap a number of transcription factor motifs.
*(Top tracks)* Gene and DMR annotation in this genomic locus.
*(Bottom right inset)* DNA methylation differences across the DMR, shown as beta-values for AF (sienna) and SR (cyan) samples. **: p-value < 0.01; ***: p-value < 0.001. Annotations below the plot correspond to individual CpGs and the sequence motifs for MNT, ARNT, SOX8, IRF6 and ZBED2 (green boxes). Sequence logos are shown for each of these transcription factors, along with expression levels of *MNT* in AF and SR samples (bottom boxplot)
*(Bottom left boxplots)* Expression levels of *SETMAR* are significantly higher in SR patients (*: p < 0.05), suggesting hypermethylation of the DMR in AF patients may reduce *SETMAR* expression. Expression levels of *SUMF1* are similar in SR and AF samples.
**b.** Gene expression levels in AF (sienna) and SR samples (cyan) for genes in the *LRRC4B* / *STY3* / *SHANK1* locus. Only *LRRC4B* expression is significantly different between control and AF patients (****: p < 0.0001; n.s.: non-significant differences). Details of annotation tracks as in Figure 7.
**c.** Cardiovascular GWAS variants over-represented in DMRs. Circle plot represents fold enrichment (circle size, log10 fold) and statistical significance (circle shade, -log10 of adjusted p-values, hypergeometric test with BH correction for multiple comparisons) for the enrichment of cardiovascular GWAS variants in all, hypermethylated or hypomethylated DMRs. GWAS variants correspond to all lead variants in the NIHR-EBI GWAS catalog plus proxy variants in strong LD for atrial fibrillation (AF), coronary artery disease (CAD), cardiomyopathies (CM) and QT interval duration (QT).
**d.** Selected transcription factor binding events overrepresented in DMRs (hyper- or hypomethylated) are represented as a circle plot. Binding data corresponds to ChIP-sequencing sets reported in cardiac tissue or stem cell-derived cardiomyocytes (iPSC-CM and ESC-CM), as catalogued in Remap 2022. Circle sizes denote the number of overlaps, and colour scale the Q-value of significance (FDR-corrected p-value from Fisher's Exact Test).
See also Tables S7 and S8.

**Table S1**: Sample and patient characteristics, including experimental readouts obtained in each sample

| Patient characteristics and clinical details | | | | | | Atrial samples used in current study | | | |
|---|---|---|---|---|---|---|---|---|---|
| **Patient ID** | **Age (yrs)** | **Gender** | **Ethnic Background** | **Disease Status** | **History of other reported conditions** | **Anatomical Side** | **H3K27ac histone modification (ChIP-seq); n = 20** | **Gene expression; n = 20** | **DNA methylation; n = 16** |
| *Discovery cohort* | | | | | | | | | |
| 25 | 80 | M | White-UK | AF | Hypertension, chronic obstructive pulmonary disease, bladder cancer | L | YES | YES | YES |
| | | | | | | R | YES | YES | YES |
| 28 | 72 | M | White-UK | AF | Hypertension, Right intracerebral bleed | L | YES | YES | YES |
| | | | | | | R | YES | YES | YES |
| 47 | 77 | M | - | AF | - | L | YES | - | YES |
| | | | | | | R | - | - | YES |
| 2231 | 84 | M | - | AF | - | L | YES | - | YES |
| | | | | | | R | YES | - | YES |
| 31 | 78 | M | White-UK | AF | Hypertension, Right Carotid Partial Blockage | L | YES | YES | - |
| | | | | | | R | YES | YES | YES |
| 313 | 58 | M | - | AF | - | L | YES | - | YES |
| | | | | | | R | YES | - | - |
| 18 | 73 | F | White-UK | AF | Hypertension, Cholycystectomy | L | - | - | YES |
| | | | | | | R | YES | - | - |
| 5 | | M | - | AF | - | L | - | YES | - |
| | | | | | | R | - | YES | - |
| 14 | | M | - | AF | - | L | - | YES | - |
| | | | | | | R | - | YES | - |
| 12b | 62 | M | White-UK | SR | Depression, bipolar disorder | L | YES | YES | YES |
| | | | | | | R | YES | YES | YES |
| 27 | 74 | M | White-UK | SR | Hypertension, Obesity | L | YES | YES | - |
| | | | | | | R | YES | YES | - |
| 45 | 91 | M | White-UK | SR | Obesity | L | YES | - | YES |
| | | | | | | R | - | - | YES |
| 20 | 59 | M | White-UK | SR | Myocardial infarction history, pulmonary tuberculosis | L | YES | YES | YES |
| | | | | | | R | YES | YES | - |
| 33 | 60 | M | White-UK | SR | Non-ST segment elevation myocardial infarction episode | L | - | - | - |
| | | | | | | R | YES | - | - |
| 7 | | M | - | SR | - | L | - | YES | - |
| | | | | | | R | - | YES | - |
| 30 | | M | - | SR | - | L | - | YES | - |
| | | | | | | R | - | YES | - |
| *Validation cohort* | | | | | | | | | |
| 9231 | 65 | M | - | AF | Hyperlipidaemia, hypertrophic cardiomyopathy | L | | | |
| | | | | | | R | | | |
| 5890 | 81 | F | - | SR | Hypertension, diabetes mellitus type I, hyperlipidaemia, chronic kidney disease, chronic obstructive pulmonary disease | L | | | |
| | | | | | | R | | | |
| 5888 | 62 | M | - | SR | Hypertension, hyperlipidaemia, severe aortic valve regurgitation | L | | N/A | |
| | | | | | | R | | | |
| 3913 | 63 | F | - | AF | Hypertension, hyperlipidaemia, chronic obstructive pulmonary disease, heart failure | L | | | |
| | | | | | | R | | | |
| 1638 | 84 | F | - | AF | Hypertension, diabetes mellitus, hyperlipidaemia, chronic obstructive pulmonary disease | L | | | |
| | | | | | | R | | | |

**Table S6.** Candidate loci selected for validation in an independent replication cohort with locus-specific quantitative PCR.

| | | Genome-wide findings (Discovery cohort) | | | | | |
|---|---|---|---|---|---|---|---|
| **Locus** | **Description** | **Gene expression** | **H3K27ac enrichment** | **Additional mechanistic evidence (external data)** | **Proposed mechanistic hypothesis** | **Potential implication in AF** | **Reference** |
| *NPPB* | Natriuretic Peptide B | AF-upregulated | Upstream AF-LA-enriched region (~14 kb) | - H3K27ac-enriched region overlapping with atrial cardiomyocyte accessible region.<br>- Enhancer region delimited by CTCF binding.<br>- H3K27ac-enriched region contacting with gene promoter.<br>- GATA4 and ESRRG binding suggests further contribution to NPPB regulation. | - Overexpression in AF resulting from increased enhancer activity. | NPPB is upregulated in response to cardiac stress, contributing to atrial remodelling through hypertrophy and fibrosis. | Tamura N, 2000; Hall E.J., 2021; Man, 2021; Sakamoto, 2022 |
| *VASH1* | Vasohibin 1 | AF-upregulated | AF-enriched regions across promoter and gene body | - H3K27ac-enriched region overlapping with atrial cardiomyocyte accessible region. | - Overexpression in AF resulting from increased promoter activity. | Inhibitor of angiogenesis with a role in the regulation of contractile kinetics of failing cardiomyocytes. It has been associated to AF, particularly in aging-related cases. | Chen C. Y., 2020; Liu C., 2023 |
| *ANGPTL2* | Angiopoietin like 2 | AF-upregulated | AF-enriched distal upstream enhancer (~60 kb) | - H3K27ac-enriched region overlapping with atrial cardiomyocyte accessible region.<br>- H3K27ac-enriched region contacting with gene promoter. | - Overexpression in AF resulting from increased enhancer activity. | Pro-fibrotic role associated with myocardial fibrosis when secreted from epicardial adipose tissue. | Kira S., 2019; Takahashi, 2022; Deniz G., 2021 |
| *STRN* | Striatin | AF-upregulated | AF-LA-enriched distal region (~65 kb) | - H3K27ac-enriched region overlapping with atrial cardiomyocyte accessible region. | - Overexpression in AF resulting from increased enhancer activity. | Cardiac striatin interacts with caveolin and calmodulin to regulate heart rate. | Nader M., 2017; Cull J.J, 2024 |
| *LRRC32* | Leucine rich repeat containing 32 | AF-upregulated | Widespread AF-LA-enriched levels across promoter and part of gene body. | - | - Overexpression in AF resulting from increased promoter activity. | Involved in formation of TGF-β, which plays a crucial role in cardiac development. Homozygous pathogenic variant reported in patient with severe dilated cardiomyopathy. | Hexner-Erlichman Z, 2022 |
| *SLC43A3* | solute carrier family 43 member 3 | AF-upregulated | AF-enriched region across promoter. | - H3K27ac-enriched region overlapping with atrial cardiomyocyte accessible region. | - Overexpression in AF resulting from increased promoter activity. | Membrane transporter protein responsible for free fatty acids uptake. Lipid metabolism dysregulation potentially contributing to AF. | Hasbargen K. B., 2020 |
| *FOXS1* | Forkhead box S1 | AF-upregulated | Widespread AF-enriched levels across gene body. | - | - Enhancer H3K27ac enriched levels contributing to upregulated expression in AF | Induced by TGF-β and found to be highly upregulated in fibrotic liver. | Bates E. A., 2024 |
| *SCX* | Scleraxis | AF-upregulated | AF-enriched promoter region | - | - Enriched H3K27ac levels within promoter contributing to increased expression in AF. | Transcription factor involved in extracellular matrix generation enabling fibroblast to myofibroblast conversion, as evidenced in in vivo deficiency models. | Bagchi RA., 2016 |
| *MT1X* | Metallothionein 1X | SR-upregulated | Widespread SR-enriched region across gene body | - H3K27ac-enriched region overlapping with atrial cardiomyocyte accessible region. | - Downregulation in AF resulting from decreased enhancer activity. | Involved in metal ion homeostasis, detoxification and protection against oxidative stress. MT1X helps degrade reactive oxygen species (ROS), which act as one of the main AF substrates. | Ding Y., 2022 |
| *RGS6* | Regulator of G protein signaling 6 | SR-upregulated | Intronic SR-RA-enriched region | - | - Downregulation in AF resulting from decreased H3K27ac levels. | Highly expressed in sinoatrial and atrioventricular nodes, where it helps controlling heart rate through inactivation of muscarinic type 2 receptor (M2Rs). | Posokhova E., 2010 |
| *NR2F1* | Nuclear receptor subfamily 2 group F member 1 | SR-LA upregulated | SR-enriched proximal enhancer (~2 kb) falling over NR2F1-AS1 anti-sense gene promoter | - H3K27ac-enriched region overlapping with atrial cardiomyocyte accessible region. | - Downregulation in AF resulting from decreased enhancer activity" | Belongs to family of orphan nuclear receptor TFs (COUP-TFs), which play a major role in atrial specification in the mouse heart. Also responsible for ion channel and electrophysiological regulation. | Devalla H. D., 2015 |
| *ANGPT1* | Angiopoietin 1 | SR-upregulated | Upstream SR-RA-enriched distal enhancer (~45 kb) | - | -Reduced H3K27ac levels in distal enhancer result in downregulated levels. | Involved in stabilization of blood vessels and plays a crucial role in vascular response to injury, suggesting a potential role in vascular inflammation in patients with AF-related stroke risk. | Kiss E. A., 2019 |
| *BLM* | BLM RecQ like heliase | SR-upregulated | SR-LA-enriched region towards 5' end | - H3K27ac-enriched region overlapping with atrial cardiomyocyte accessible region.<br>- H3K27ac-enriched region contacting with gene promoter. | - Reduced H3K27ac levels towards 5' end of gene result in downregulated levels. | Crucial role in DNA repair processes and genomic stability maintenance. Loss of DNA repair can increase oxidative stress levels and deteriorates overall cardiac function. | Kaur E., 2021 |
